# Calcium-Based Synaptic and Structural Plasticity Link Pathological Activity to Synaptic Reorganization in Parkinson’s Disease

**DOI:** 10.1101/2025.03.06.641850

**Authors:** Cathal McLoughlin, Justus A. Kromer, Madeleine Lowery, Peter A. Tass

**Affiliations:** Department of Electrical and Electronic Engineering, University College Dublin, Dublin & D04 V1W8, Ireland; Department of Neurosurgery, Stanford University, Stanford & California, United States of America

## Abstract

Altered motor symptoms of Parkinson’s disease (PD) are associated with dopaminergic neuronal loss. Widespread synaptic reorganization and neural activity changes, including exaggerated beta oscillations and bursting, follow dopamine depletion (DD) of the basal ganglia (BG). Our computational model examines DD-induced neural activity changes and synaptic reorganization. It encompasses the BG sub-circuit comprising the subthalamic nucleus and globus pallidus externus. Calcium-dependent synaptic and structural plasticity mechanisms were incorporated, allowing neural activity to alter network topology. We show how elevated iMSN firing rates can induce synaptic connectivity changes consistent with PD animal models. We suggest synaptic reorganization following DD results from a series of homeostatic calcium-based synaptic changes triggered by elevated iMSN activity. Structural plasticity counteracts DD-induced neural activity changes and opposes exaggerated beta oscillations, whereas synaptic plasticity alone amplifies beta oscillations. Our results suggest that synaptic and structural plasticity have qualitatively different contributions to DD-induced synaptic reorganization in the BG.

**Teaser:** Synaptic and structural plasticity differentially drive synaptic reorganization and abnormal activity in Parkinson’s disease.

## Introduction

### Symptoms, Causes, and Pathways of Parkinson’s Disease

Parkinson’s disease (PD) is a slowly progressing neurodegenerative disorder characterized by both motor symptoms (such as tremors, rigidity, akinesia, and bradykinesia) and non-motor symptoms (such as apathy and sleep problems (*1*)). As the second most common neurodegenerative disorder, PD affects approximately 2-3% of people 65 years and older (*2*).

PD motor symptoms appear following the loss of midbrain dopaminergic neurons, resulting in dopamine depletion (DD) in the basal ganglia (BG); however, the exact mechanisms that lead to PD motor symptoms are still undetermined (*3*). In the late 1980s, Albin et al. (*4*) and DeLong (*5*) hypothesized a direct/indirect pathway model of BG functionality. The model suggested that BG disorders, including PD, result from an imbalance between the prokinetic direct and the akinetic indirect pathways. The loss of dopaminergic striatal inputs differentially affects indirect pathway (iMSNs) and direct pathway striatal medium spiny projection neurons (dMSNs), leading to the dominance of the indirect over the direct pathway (*6*). This results in elevated iMSN activity, increased inhibition of the external globus pallidus (GPe), and disinhibition of the subthalamic nucleus (STN) (*7*). Ultimately, this increases the firing rates of BG output neurons, e.g., in the internal globus pallidus (GPi), and over-inhibits thalamocortical motor circuits (*4, 5*) (Figure 1a,b). Empirical tests of the direct/indirect pathway model in animal models of PD confirmed that bilateral excitation of iMSNs elicited hypokinetic motor symptoms such as freezing and bradykinesia, stressing the critical role of elevated iMSN activity in PD. In contrast, excitation of dMSNs reduced freezing and increased locomotion (*9*). Despite these observations, the delineation of striatal outputs into orthogonal direct and indirect pathways was shown to be less distinct than initially supposed (*6*). Furthermore, the relationship between BG output activity and motor symptoms is inconsistent (*10, 11*).

**Figure 1:**
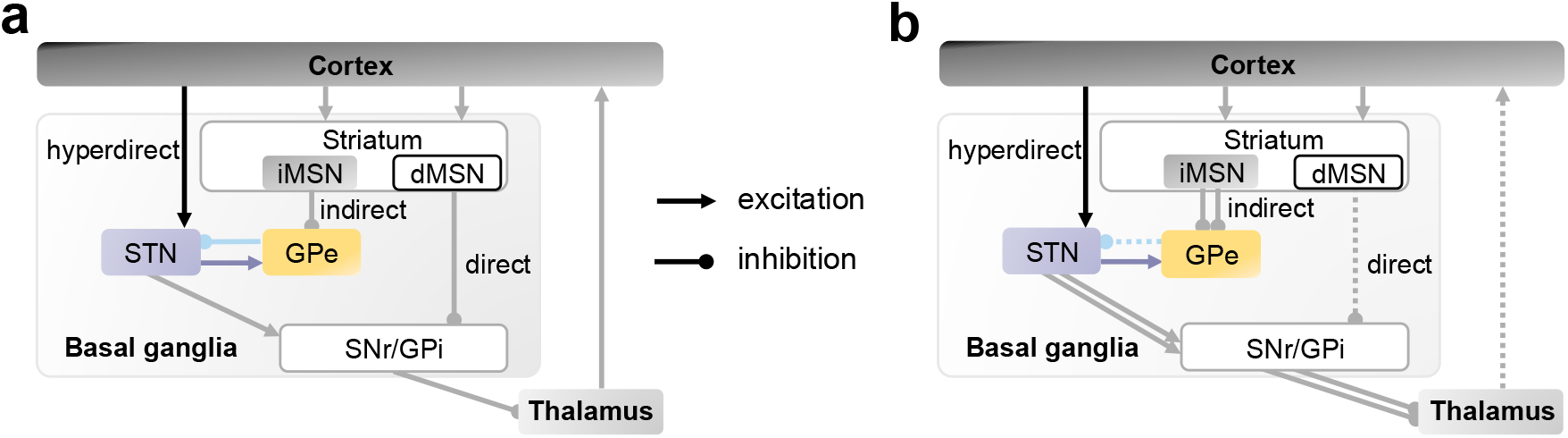
Schematic diagram of synaptic connectivity in the basal ganglia (BG). **a:** According to the direct/indirect pathway model of PD, dopamine depletion leads to an imbalance between the direct cortex-dMSN-substantia nigra pars reticulata (SNr)/GPi-thalamus pathway and the indirect cortex-iMSN-GPe-STN-SNr/GPi-thalamus pathway (*4, 5*). Later, the akinetic hyperdirect pathway was identified (*8*). Intra-nucleus connections are not shown. b: Changes in BG network in PD according to direct/indirect pathway model (*4, 5*). Dotted arrows represent weakening, and doubled arrows strengthening, relative to the healthy state. Note that STN→GPe and direct CTX→STN connections were not included in the original model (*4, 5*).

### The Role of Beta Oscillations in the STN-GPe Circuit

Later, it was suggested that pathological activity patterns, such as bursting and excessive synchronous oscillations, underlie PD motor symptoms by preventing individual neurons from independently processing motor-related information (*3, 12*). However, conflicting observations exist regarding the emergence and outcome of varying beta oscillations. Some earlier experimental evidence supported the pathological role of beta oscillations in causing hypokinetic symptoms (*13–15*); however, this was later challenged by the observation of excessive beta oscillations in both PD and dystonia patients, as the latter typically suffer from hyperkinetic symptoms (*16*). Furthermore, hypokinetic motor symptoms such as akinesia precede the onset of beta oscillations in animal models of PD (*17–19*). In other published studies using rodent models of PD, it was also observed that chronic, but not acute, DD leads to exaggerated beta oscillations (*17, 19*).

Therefore, given these conflicting results, we focused on examining beta oscillations within the STN-GPe circuitry. The STN-GPe circuit lies at the convergence of the hyperdirect (*8*) and indirect pathways (Fig. 1); it is a crucial area where beta oscillations manifest in PD (*20*). Experimental studies in the non-human primate model for PD suggest a vital contribution of the STN to excessive beta oscillations (*21*). Modeling studies have further analyzed the ability of the STN-GPe circuit to generate and maintain beta oscillations (*22, 23*). Recent studies in rats reported that cortical and GPe beta oscillations persist after optogenetic STN inhibition, whereas GPe inhibition suppresses such oscillations. The authors concluded that in rats, the GPe has a pivotal role in generating beta oscillations (*24*). Later computational studies pointed out that different circuits might be involved in generating beta oscillations in various animal models and PD patients (*25*).

Given that beta oscillations in animal models of PD emerge after early motor symptoms, it has been suggested that they appear because of slow adaptation processes in related brain networks (*17–19, 26*). Synaptic reorganization, including changes in synaptic strengths and/or numbers, has been proposed as a mechanism underlying such processes (*26*). Recent experiments in rodent models of PD reported substantial synaptic reorganization in the BG following DD. DD was shown to change the strength of synapses (synaptic plasticity) and the number of synaptic connections (structural plasticity) between several populations of BG neurons (*27–31*). Known synaptic changes in the STN-GPe circuit encompass both excitatory and inhibitory synaptic connections (Fig. 2) and likely impact network dynamics; that is, their ability to generate oscillatory activity (*26*).

**Figure 2:**
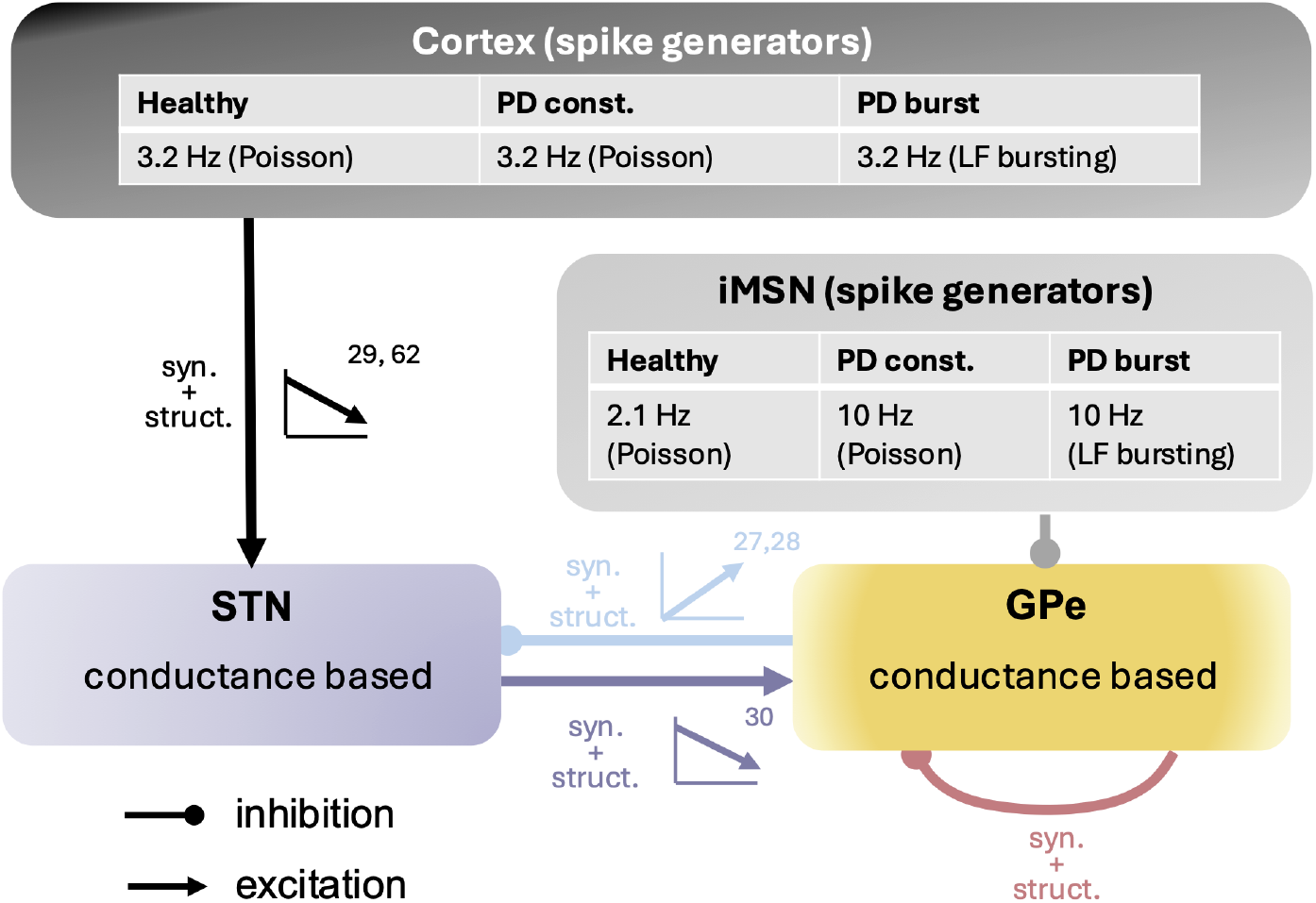
Schematic diagram of our computational model of the STN-GPe circuit. Cortical and iMSN inputs were approximated by Poisson spike generators, with a time-dependent spike rate adjusted to mimic the healthy state, and two PD states: one with fixed cortical and iMSN Poisson rate parameters (PD const.) and one with time-dependent cortical and iMSN Poisson rates mimicking low-frequency bursting (PD burst.) (see Methods for details). STN and GPe neurons were described by conductance-based point neuron models previously published in Refs. (*32*) and (*33*). Labels next to synaptic connections indicate which types of plasticity were considered with “syn.” referring to calcium-based synaptic plasticity implemented according to Graupner and Brunel (*34*) and “struct.” structural plasticity, which we modeled using a linearized version of the Butz and Van Ooyen framework (*35*). Small arrows indicate whether an increase or a decrease in overall synaptic inputs were observed in PD animal models, respectively (see corresponding references).

Whether and how these synaptic changes relate to observed BG activity changes, such as the emergence of beta oscillations, remains unclear. STN-GPe reorganization is observed via an increase in the number and strength of GPe synapses onto STN neurons (*27, 28*), the depression of cortico-STN (*29*) and STN-GPe excitatory inputs (*30*), and other structural changes within the striatum. Mallet et al. suggested that this reorganization might represent a homeostatic change that counteracts elevated input from iMSNs and resulting activity changes in the indirect pathway, as described in the direct/indirect pathway model (*26*). As it is likely that synaptic reorganization acts on slower time scales than DD-induced striatal activity changes, it might also contribute to the late emergence of pathological beta oscillations (*17–19*).

### Current Treatments for Akinetic Symptoms of PD

The ability to induce positive therapeutic effects that outlast the stimulation period may reduce the overall required stimulation power and the risk of unwanted side effects from current treatments such as high-frequency DBS (*36*). Coordinated reset stimulation (CRS) (*37*) is a promising computationally developed technique that extends therapeutic effects beyond the initial stimulation. Studies conducted using neuronal network models incorporating synaptic plasticity predicted stimulation-outlasting effects (*38*). In plastic networks, stable synchronized states (mimicking synchronized pathological oscillations in PD) and stable desynchronized states (mimicking the absence of synchronized pathological oscillations) can coexist. During CRS, synaptic connectivity changed, which drove the networks from a synchronized state into a stable desynchronized state, inducing stimulation-outlasting desynchronization effects (*39–41*). Preclinical studies in the non-human primate PD model showed that CRS of the STN induced corresponding cumulative and long-lasting desynchronization and therapeutic effects that outlasted stimulation by several weeks (*39, 42, 43*). CRS was successfully delivered to PD patients using DBS electrodes (*44*) and vibrotactile fingertip stimulation (*45, 46*). However, only recently has CRS-induced structural plasticity been studied computationally in networks of phase oscillators (*47*) and the STN-GPe circuit, where excitatory connections were plastic and synaptic and structural plasticity were not simulated together (*48*). Recently, studies in DD mice also reported stimulation-outlasting therapeutic effects after inhibition of one type of GPe neurons (*49*). Based on these results and the observation that different types of GPe neurons responded in opposite ways to the onset of high-frequency electrical stimulation, Spix et al. developed a DBS-burst stimulation technique that led to cell-specific responses in brain slices and to stimulation-outlasting therapeutic effects in DD mice (*50*). The ability to model synaptic reorganization in the STN-GPe circuit during stimulation targeting stimulation-outlasting and cumulative effects, such as CRS and pallidal burst stimulation (*51*) may help develop clinically testable hypotheses about effective stimulation parameters as well as underlying mechanisms.

### A Computational Model to Examine Synaptic Reorganization in the STN-GPe Circuit

We suggest that synaptic reorganization following DD results from a series of homeostatic calcium-based synaptic changes triggered by elevated iMSN activity. The critical role of elevated iMSN activity in PD pathology presents a core tenet of the direct/indirect pathway model and has been shown experimentally (*9*). However, a model of long-term synaptic plasticity in the BG that explains the observed synaptic changes after chronic dopamine depletion has not been presented to date. A computational model, in which the relationship between observed behavior and neuronal properties can be directly compared, may help develop future therapeutic strategies that induce synaptic reorganization to promote long-term effects.

We developed a computational model of calcium-based synaptic and structural plasticity in the STN-GPe circuit, a part of the BG that is believed to be involved in the generation and/or maintenance of exaggerated beta oscillations in PD (*21, 22, 33*) (Fig. 2). In particular, we model the calcium concentrations in local dendritic segments and their modulations due to pre- and postsynaptic activity. To achieve the performance necessary for multi-day simulations with sub-millisecond integration timesteps, we implemented the model using the GPU parallelization library CUDA.

Our model not only reproduces currently available data on long-term plasticity in the STN-GPe circuit but also demonstrates that synaptic and structural plasticity may contribute to the observed synaptic reorganization presenting homeostatic adaptation to counteract increased indirect pathway activity according to the direct/indirect pathway model (*4,5*). Our results show that an increase in the firing rate of iMSNs is sufficient to trigger structural synaptic changes in the STN-GPe sequence. We further observe distinct calcium-based structural and synaptic plasticity contributions in response to beta activity elevation.

We combine existing computational models of STN and GPe neurons with detailed computational models of calcium-based synaptic and structural plasticity to better investigate the relationship between BG neural activity patterns and synaptic reorganization. By doing so, we hope to facilitate the refinement of current therapies and the development of novel approaches to treating Parkinson’s disease.

## Results

### Development of the Computational Model

We developed a computational model of long-term plasticity in the STN-GPe circuit consisting of 250 STN and 750 prototypic GPe neurons simulated using modified conductance-based models (*32, 33*). Cortical and iMSN activities were modeled by Poisson spike generators. Synapses were represented using the multi-state model Rubin et al. (*52*). Excitatory synapses contain AMPA and N-methyl-D-aspartate (NMDA) receptors, while inhibitory synapses contain GABAa receptors. Acute changes in calcium concentrations alter synaptic conductances, which we model using the framework from Graupner and Brunel (*34*). Their model utilizes differing calcium concentration thresholds; below a minimum threshold, there is no change in synaptic efficacy, whereas above this threshold, synaptic depression occurs. At high calcium concentrations above an upper threshold, synaptic potentiation occurs.

Additionally, we implement homeostatic structural plasticity to maintain an average neural calcium concentration, following the work of Butz and Van Ooyen (*35*). Striatal and cortical inputs are simulated with Poisson spike generators, and their activity is manipulated to study synaptic reorganization in response to changes in striatal and cortical inputs mimicking those observed in PD.

The nature of plasticity at synapses onto STN neurons has been investigated experimentally (*28, 53*). Plasticity at CTX to STN synapses is dependent on pre- and postsynaptic activation. Optogenetic stimulation of cortical axons can induce long-term potentiation (LTP) of CTX to STN synapses; LTP is mediated by calcium influx into postsynaptic STN neurons via NMDA receptors (*28*). Interestingly, GPe to STN synapses are potentiated concomitantly, demonstrating that plasticity in the STN is heterosynaptic (*54*). Co-potentiation of CTX to STN and GPe to STN synapses appears to balance the levels of excitation and inhibition within the STN neurons (*28*) and thus may act to maintain STN activity at some physiological set point.

In its default state, the model was tuned to reflect the dopamine-intact STN-GPe circuit of the rat, for which the neural firing patterns and rates are well-described in the literature (*55, 56*).

The results demonstrate that our computational model reproduces known characteristics of synaptic plasticity in the STN-GPe circuit, including heterosynaptic plasticity in the STN (*28, 53*) and long-term plasticity induced by rebound bursting (*53*).

### Heterosynaptic Plasticity in the STN

Heterosynaptic plasticity was modeled by specifying that each neuron has three dendritic segments, each containing local calcium concentrations that influence the strengths of all synapses in that segment (Fig. 6).

To test whether our model can reproduce the dynamics of cortically evoked synaptic plasticity in the STN, we established an experimental setup similar to that of Chu et al. (*28*); cortical stimulation was delivered, and all synaptic inputs were blocked, except for CTX to STN NMDA receptors (see Fig. 3a). This protocol was capable of inducing LTP of STN-bound synapses in our model. A representative STN neuron’s response to corresponding cortical input is shown in Fig. 3b. Calcium influx occurs through NMDA receptors when the time delay between pre- and post-synaptic spiking is short, increasing the local calcium concentrations in the dendritic segments (Fig. 3b). As both excitatory and inhibitory synapses may be bound to the same dendritic segment (*57*), acute changes in local calcium concentrations lead to co-potentiation of both synapse types. Repeated exposure to cortical stimulation resulted in acute changes in STN calcium concentrations. This led to LTP in CTX to STN and GPe to STN synapses (Fig 3c and d), reproducing the effect of heterosynaptic plasticity (*28*).

**Table 1:** STN and GPe ionic currents. Dashed horizontal line separates STN-only currents (below) from STN and GPe ionic currents (above).

| Symbol | Name | Equation |
| --- | --- | --- |
| $I_i^{\text{Na}}$ | Sodium current | $g^{\text{Na}} m_i^3 h_i (V_i - E^{\text{Na}})$ |
| $I_i^{\text{K}}$ | Kv3-type potassium current | $g^{\text{K}} n_i^4 (V_i - E^{\text{K}})$ |
| $I_i^{\text{T}}$ | T-type calcium channel | $g^{\text{T}} p_i^2 q_i (V_i - E_i^{\text{Ca}})$ |
| $I_i^{\text{Ca-K}}$ | Calcium-activated potassium current | $g^{\text{Ca-K}} r_i^2 (V_i - E^{\text{K}})$ |
| $I_i^{\text{leak}}$ | Leak current | $g^{\text{leak}} (V_i - E^{\text{leak}})$ |
| $I_i^{\text{A}}$ | Voltage-dependent A-type potassium current | $g^{\text{A}} a_i^2 (V_i - E^{\text{K}})$ |
| $I_i^{\text{L}}$ | L-type long-lasting calcium current | $g^{\text{L}} c_i^2 d_{1i} d_{2i} (V_i - E_i^{\text{Ca}})$ |

**Figure 3:**
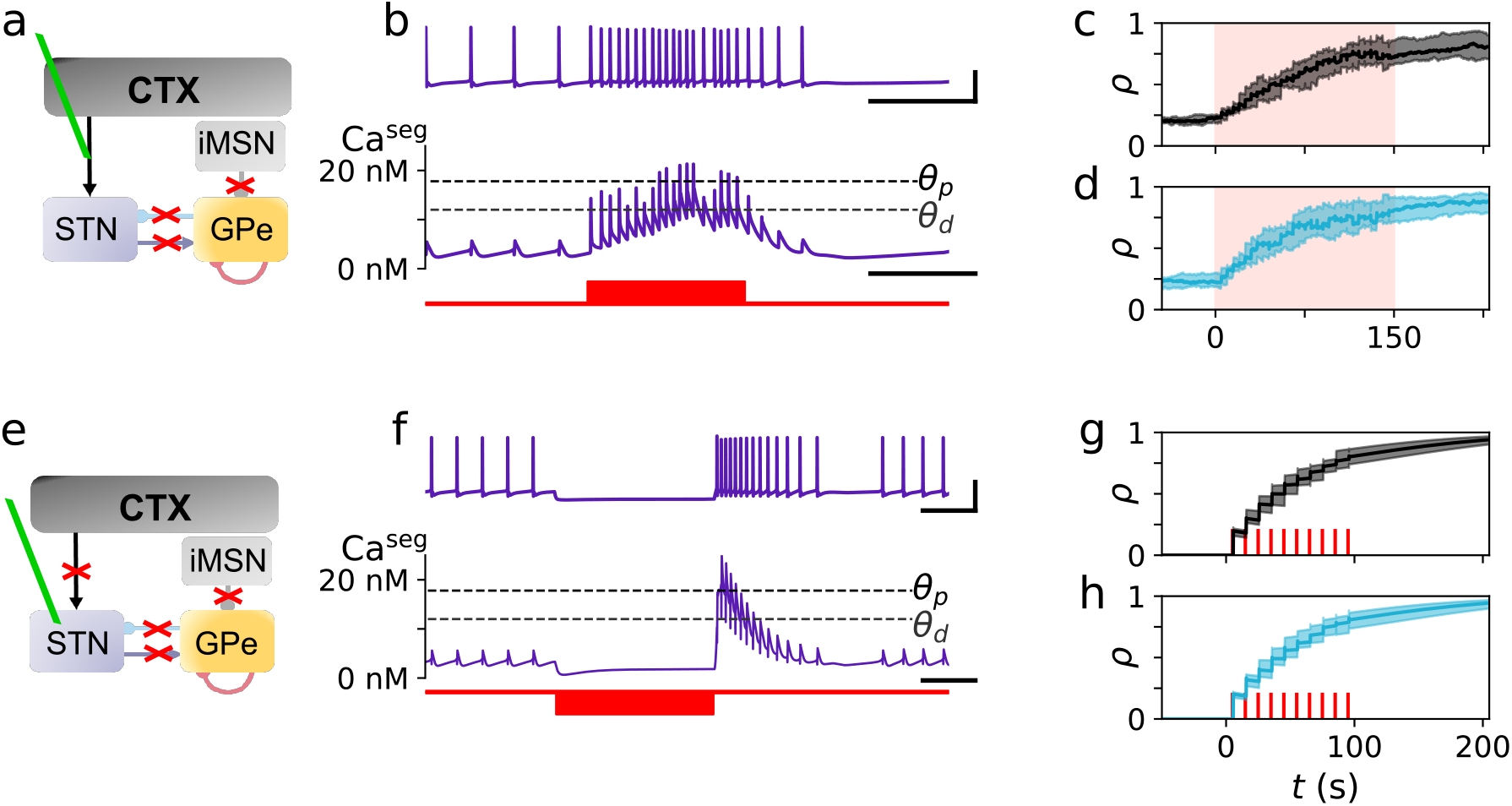
Reproducing plasticity experiments. **a:** Model layout for reproducing the results from Chu et al. (*28*) where all non-NMDA related synaptic currents onto the STN are blocked (red crosses). The green line illustrates that stimulation is applied at the CTX to STN connections. b: STN activity (b, top), Ca concentration within a dendritic segment, Ca^seg^, (b, center), and cortical stimulation, implemented by increasing the firing rate of Poisson cortical inputs to the STN to 50 Hz for 300 ms every 5 sec (b, bottom). Horizontal dashed lines mark the depression threshold, Θ _d_, and potentiation threshold, Θ _p_, of the synaptic plasticity model (*34*). c,d: LTP resulting in an increase of the synaptic efficacy, d, is observed at CTX to STN (c) and GPe to STN (d) synapses following 150 s of cortical stimulation (red). e: Model layout for reproducing the stimulation protocol from Wang et al. (*53*). f: STN membrane potential (f, top), Ca^seg^, (f, center), during a rebound burst induced by a hyperpolarizing current pulse of −10 ‘A/cm^2^ (f, bottom). Black horizontal bars in b and f mark time intervals of 200 ms, and vertical bars mark a voltage difference of 50 mV. g,h: Repeated rebound bursts induced by a sequence of 10 current pulses delivered to the STN at a rate of 0.1 Hz lead to LTP of CTX to STN (g) and GPe to STN (h) synapses. c,d,g,h: The median and interquartile range of results from ten simulations are shown.

### Rebound Bursting-Induced Long-Term Plasticity

In their experiments, Wang et al. (*53*) found that the strength of GPe to STN synapses could be altered via rebound bursting of STN neurons induced by hyperpolarizing current pulses. The number of spikes in a rebound burst determined whether no synaptic changes, long-term depression (LTD), or LTP occurred. They linked these effects to the quantity of calcium influx by systematically blocking T- and L-type calcium channels (*53*).

To demonstrate that our computational model can reproduce the dynamics of inhibition-induced long-term plasticity, we established an experimental setup similar to theirs (see Fig. 3e) (*53*). Synaptic transmission was blocked, and hyperpolarizing current pulses were delivered.

A representative response of an STN neuron to a single current pulse is shown in Fig. 3f. Hyperpolarization de-inactivates T-type calcium channels, facilitating calcium influx during the rebound burst (*58*) (Fig. 3f). As in Wang et al., short bursts did not change synaptic efficacy (*53*), as calcium remained below the depression threshold. Short bursts resulted in LTD when this threshold was crossed. Long bursts resulted in potentiation as calcium crossed the potentiation threshold (Fig. 3f). Repeated exposure to sufficiently long rebound bursts led to potentiation of GPe to STN synapses (Fig. 3h), similar to the results of Wang et al. (*53*).

These results support the hypothesis that homo- and heterosynaptic plasticity in the STN can be explained by a calcium-based synaptic plasticity mechanism similar to the one presented by Graupner and Brunel (*34*). Due to the lack of experimental evidence, we assumed that a similar plasticity model applies to the GPe, as heterosynaptic plasticity is seen in many diverse types of neurons (*54*). Specifically, we assumed the same mechanism is present at all synapses except for the iMSN to GPe synapses (*59*).

### Increased Indirect Pathway Activity Triggers Homeostatic Synaptic Reorganization in the STN-GPe Circuit

In PD, the loss of striatal dopamine promotes hyperactivity of iMSNs (*4–6*). As hypothesized by previous studies, compensatory synaptic reorganization in the BG might occur to counteract elevated iMSN activity (*26*). To test this hypothesis, we studied synaptic reorganization in response to an instantaneous increase in the iMSN firing rate.

Our computational model incorporated homeostatic, calcium-based, activity-dependent structural plasticity based on the linear Butz and Van Ooyen algorithm (*35, 60*). With this mechanism, the number of excitatory and inhibitory connections at each neuron changes in response to deviations of low-pass-filtered somatic calcium concentrations from a target calcium concentration. The timescale at which synaptic connections change is of the order of hours. Updates to synaptic connectivity occur every 200 ms. Each excitatory and inhibitory connection is associated with a randomly chosen segment on the post-synaptic neuron.

We first simulated networks for 13 hours to obtain stationary states. Ten stationary states were obtained from simulations beginning with random initial conditions for neuron and synapse gating variables and random connections between neurons (Methods). We performed simulations from each of these states, where the firing rate of the iMSNs was increased from 2.1 Hz to 10.0 Hz (Fig. 4b); we recorded firing rates and calcium concentrations of each neuron, synaptic efficacies, and in-degrees of each synapse type. In the following text, we refer to the mean of these data.

**Figure 4:**
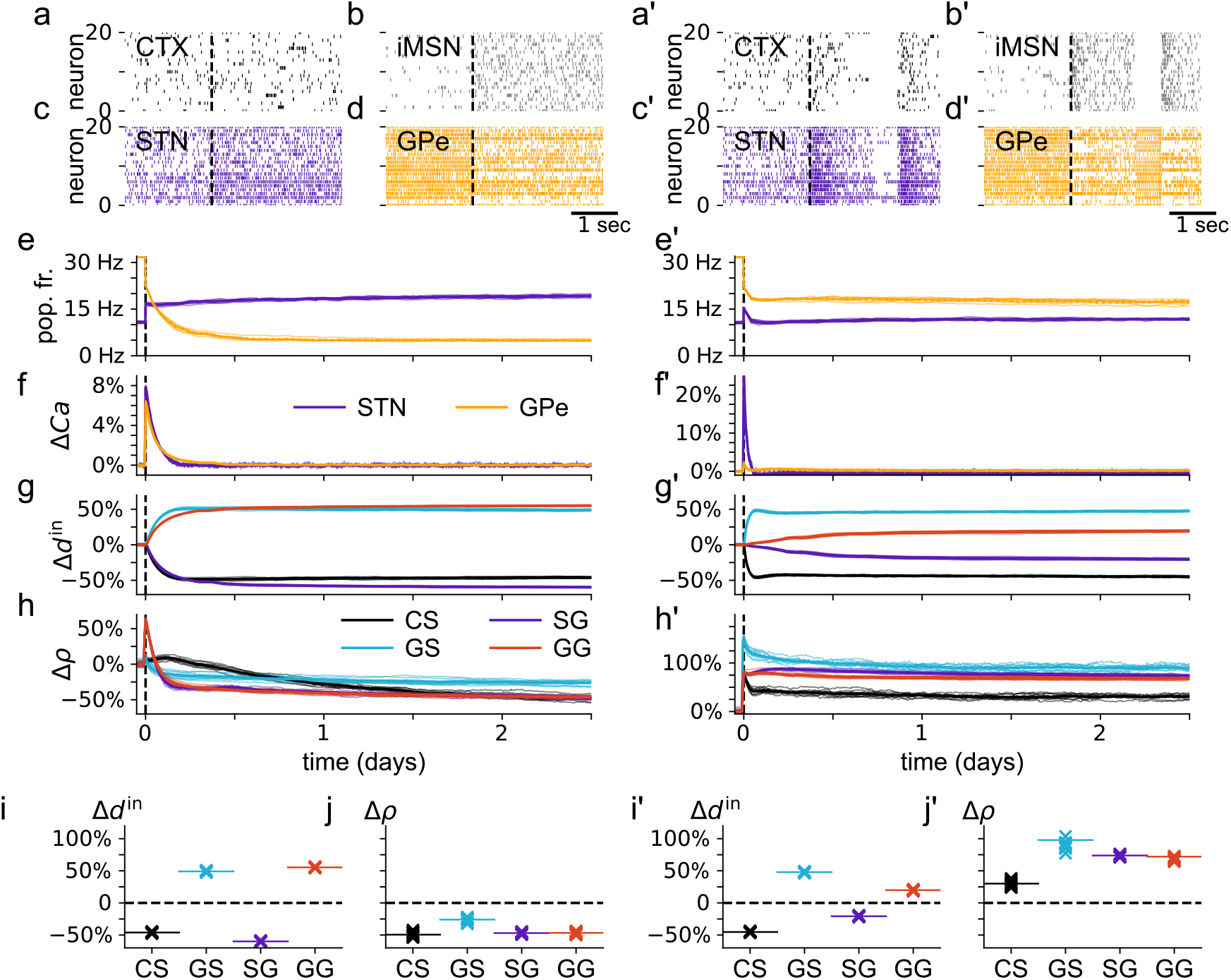
Instantaneous changes of iMSN and cortical inputs induce synaptic reorganization. **a-j:** iMSN firing rate was increased from 2.1 Hz to 10.0 Hz at t = 0 hours while the cortical rate was kept unchanged (vertical dashed black lines in a-h). a,b,c,d: Raster plots before and after iMSN firing rate change. For each curve in e,f,g, and h, thin lines correspond to traces from individual simulations, while thick lines correspond to the mean of 10 simulations. e: Mean instantaneous firing rates in the STN and GPe throughout the simulation. f: Change of population-averaged low-pass-filtered somatic calcium concentration in STN and GPe relative to its value before the iMSN firing rate change (preIMSN). g: Change of average in-degree of different synapse types relative to in-degree preIMSN. h: Average synaptic efficacies of each synapse type relative to preIMSN values. Final values are shown in panels i (average in-degrees) and j (average synaptic efficacies) (see Methods for details). We used the abbreviations CS for CTX to STN, GS for GPe to STN, SG for STN to GPe, and GG for GPe to GPe synapses. a’-j’: Same as a-j, but cortical and iMSN inputs were instantaneously switched to low-frequency bursting (0.5 Hz) at t=0. The overall mean instantaneous firing rate of cortical and iMSNs after the activity change was 3.2 Hz (a’) and 10 Hz (b’), respectively (see Methods for details on bursting activity).

The firing rate of STN neurons increased, and the firing rate of GPe neurons decreased in response to increased iMSN activity (Fig. 4c, 4d, and 4e). This further caused transient elevations in somatic calcium concentrations in both the STN and GPe (Fig. 4f). Synaptic reorganization was triggered as a result, including changes in the mean number of incoming connections to a neuron (in-degrees) (structural plasticity, Fig. 4g) and changes in synaptic efficacies (synaptic plasticity, Fig. 4h). We observed transient synaptic dynamics on the time scale, between several hours and 1.5 days, after which synaptic connectivity stabilized.

All synaptic efficacies were transiently enhanced but declined below their initial value by the end of the simulation (Fig. 4h). In addition to weakening individual synapses, structural plasticity promoted lower in-degrees for CTX to STN and STN to GPe connections and higher in-degrees for GPe to STN and GPe to GPe connections, corresponding to losses and gains of respective synapse types (Fig. 4g).

These results support the hypothesis that elevated iMSN activity, which is at the core of the indirect/pathway model, triggers synaptic reorganization in the indirect pathway via synaptic and structural plasticity.

### Low-Frequency Bursts Strengthen Synaptic Connections

We have shown that the iMSN firing rate increase alone is sufficient to alter synaptic connectivity in the STN-GPe circuit. Next, we tested how low-frequency burst inputs—commonly observed in PD patients and related animal models (*56, 61*)—affect synaptic connectivity. Ten simulations were performed after establishing stationary states, as described above; we instantaneously changed the activity of cortical and iMSN Poisson spike generators to a low-frequency burst pattern while keeping the same mean firing rates as in the previous section, resembling experimentally observed bursting activity in PD (*56, 61*).

As shown in Fig. 4c’ and 4e’, the mean firing rate of STN neurons increased slightly, and that of GPe neurons decreased (Fig. 4d’) following the cortical and iMSN activity change. However, GPe firing rates were higher than after an instantaneous increase of the iMSN firing rate (compare Figs. 4e and 4e’). The transient change in the population-averaged low-pass-filtered somatic calcium concentration in the STN shown in Fig. 4f’ was noticeably larger than in the previous section (Fig 4f). In contrast, the change in GPe calcium concentration was lower (compare Figs. 4e’ and 4e).

Changes in in-degrees were qualitatively similar to those observed following the iMSN firing rate increase (compare Figs. 4i’ and 4i); however, the changes in STN to GPe and GPe to GPe in-degrees (Fig. 4g’) were weaker and occurred on a slower time scale. In contrast to the iMSN rate increase, cortical and iMSN low-frequency bursting resulted in persistent increases in synaptic efficacies in the STN-GPe circuit (Fig. 4h’).

Our computational results suggest that both scenarios increased (constant) iMSN firing rates and low-frequency bursting (with the same mean firing rate of iMSNs) led to qualitatively similar changes in the number of synapses (structural plasticity). In contrast, while an increase in the iMSN firing rate resulted in weaker synapses in the STN-GPe circuit (Fig. 4j), cortical and iMSN low-frequency burst input led to stronger synapses (synaptic plasticity) (Fig. 4j’).

### Synaptic Plasticity Shapes the Evolution of Beta Oscillations in the STN-GPe Circuit

Exaggerated beta oscillations are associated with akinetic motor symptoms, such as bradykinesia and rigidity, that are observed in recordings from human patients (*62*) and animal models (*17*) of PD. Computational models suggest that alterations in neural connectivity can alter the level of synchronous beta oscillations (*25, 63*). Therefore, we examined the time course and beta oscillation activity level in the STN and GPe populations following changes in iMSN and/or cortical activity discussed in the previous sections (Fig. 5). The following data were derived from 10 simulations, which continued from the stationary states established in our previous numerical experiments.

**Figure 5:**
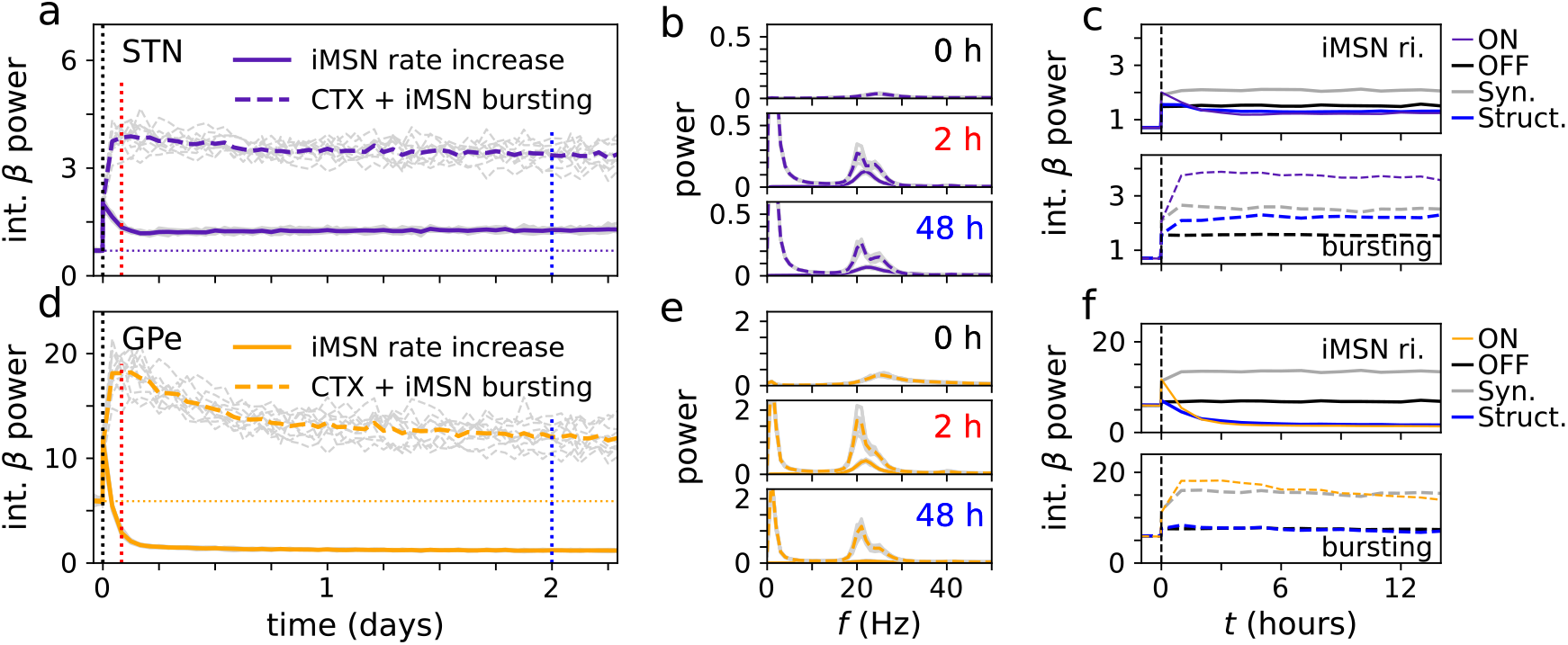
Evolution of beta power following instantaneous changes of input activity. Ten trials were simulated, including different random initial conditions and noise realizations. Colored lines mark averages over trials, and gray lines results for individual trials. a: Computational results showing the integral STN V power (integral power over a frequency range of 13-30 Hz) for simulations of network response to instantaneous iMSN rate change (solid curves) and instantaneous change of CTX-iMSN burst input (dashed curves). The horizontal dotted line marks pre-input activity change conditions. b: Corresponding power spectra at different times after activity change (labels and accordingly colored vertical lines in a). c: Comparison of trial-averaged integral beta power for simulations with synaptic and structural plasticity (ON), to simulations without plasticity (OFF), and to simulations in which either only synaptic plasticity was switched on (Syn.) or only structural plasticity was switched on (Struct.) after input activity change. d-f: Same as a-c, but for GPe activity.

**Figure 6:**
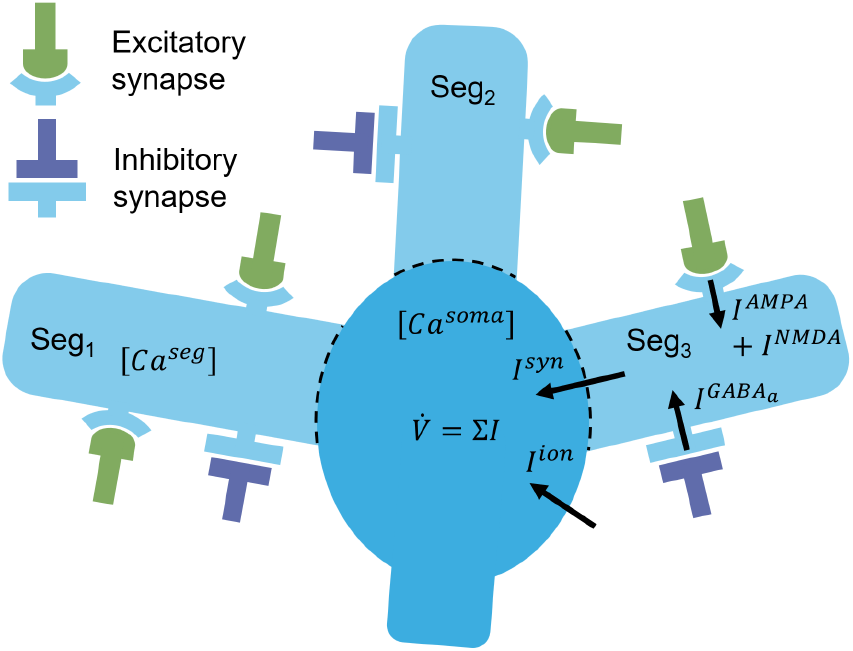
Schematic diagram of a single model neuron in this study. Each neuron contains three isolated segments (Seg_1_, Seg_2_, Seg_3_), where local calcium concentrations (Ca^seg^) are modeled. These local calcium concentrations influence the efficacy of the excitatory and inhibitory synapses that are present on the segment. Currents contributing to changes in membrane potential, *V*_i_, arise from: voltage-gated ion channels, *I*^ion^ (Table 1), excitatory (*I*^NMDA^ and *I*^AMPA^), and inhibitory (*I*_GABA_) synapses (*I*^syn^). Currents contributing to changes in segmental and somatic calcium concentrations arise from T-(*I*_T_) and L-(*I*_L_) type calcium channels, and from the calcium component of NMDA receptors on excitatory synapses (*I*_NMDA,Ca_). Segment calcium concentrations are influenced by their local NMDA currents, whereas somatic calcium concentrations are influenced by the NMDA currents from all synapses on the neuron.

Both the increase in the iMSN firing rate and low-frequency bursting led to a rapid increase in STN and GPe beta power, which lasted several hours. Afterward, beta power stabilized at higher values than before the change, except the GPe power, which stabilized at lower values after the increase in the iMSN firing rate. In our computational model, it took about 12 hours to a day for the beta power to stabilize.

Next, we explored the contributions of both synaptic and structural plasticity to the emergence of beta oscillations following sudden changes in iMSN and/or cortical inputs. We performed simulations in which either synaptic plasticity, structural plasticity, or both types of plasticity were switched off. Corresponding results are shown in Fig. 5c and 5f. Simulations with synaptic plasticity displayed higher beta power than those without synaptic plasticity, whereas simulations with structural plasticity tended to have lower beta power than those without structural plasticity. An exception to the latter is the STN beta power response to cortical and iMSN low-frequency bursting in which structural plasticity also elevated the beta power relative to simulations without structural plasticity (Fig. 5c). We also observed an increase in STN and GPe beta power in response to activity changes in the absence of synaptic and structural plasticity (Fig. 5c,f).

These results suggest that synaptic plasticity, in response to changes in iMSN and/or cortical input activity, increases STN and GPe beta power. In contrast, slower synaptic reorganization due to structural plasticity counteracts elevated beta oscillations in the GPe and STN during constant iMSN firing rate increases. Due to the elevation in beta power observed in the absence of synaptic reorganization (Fig. 5c,f), our results also suggest that a substantial portion of the beta power change is due to upstream changes of neuronal input activity in the indirect and/or hyper-direct pathway.

## Discussion

### Synaptic Reorganization as a Compensatory Mechanism

PD is accompanied by widespread activity changes and synaptic reorganization in the BG. Historically, models of BG dysfunction have suggested that increased iMSN activity (*4, 5*), and/or exaggerated beta oscillations, contribute to akinetic PD motor symptoms such as rigidity and bradykinesia (*62*). However, the ways in which BG activity patterns and synaptic reorganization relate to each other, and ultimately contribute to PD motor symptoms, are still in the process of being understood.

Previous studies suggested that some synaptic reorganization occurs as a compensatory mechanism to counteract neuronal activity changes in the indirect pathway that follows iMSN hyperactivity. In particular, the increase in the number and strength of GP to STN synapses and the depression of cortical to STN inputs (*25, 26*). Motivated by this hypothesis, we explored the role of different plasticity mechanisms in synaptic reorganization following iMSN hyperactivity. To this end, we build a detailed computational model of long-term synaptic and structural plasticity in the STN-GPe circuit, a major part of the indirect pathway.

We incorporated the calcium-based synaptic plasticity mechanism presented by Graupner and Brunel (*34*) into a network model of the STN-GPe circuit, which contains conductance-based neuron and synapse models. In the model, two thresholds separate calcium concentrations for which either no synaptic change, synaptic depression, or synaptic potentiation occurs. We showed that such calcium-based synaptic plasticity reproduces previous experimental results on hetero- and monosynaptic plasticity in rodent STN (*28, 53*).

In our computational model, repeated bouts of elevated cortical inputs induced STN bursting and led to heterosynaptic plasticity of CTX to STN and GPe to STN synapses (Fig. 3). NMDA receptor-mediated increases in STN segment calcium concentrations caused the latter. This agrees with previous experiments on heterosynaptic plasticity in mice models, namely, in which LTP of CTX to STN and GPe to STN synapses was observed when pre-synaptic cortical axons and post-synaptic STN neurons were activated simultaneously (*28*). In experimental rat models, Wang et al. studied monosynaptic plasticity in the STN, linking the strength of hyperpolarization-induced rebound bursting to changes in the strength of GPe to STN connections (*53*). Very weak rebound bursts induced minor to no changes in synaptic strength, weak bursts caused LTD, and strong bursts induced LTP (*53*). Mirroring their animal modeling in our computational model, we found that corresponding synaptic long-term changes occurred if rebound bursting-induced calcium changes led to no threshold crossing, crossing of the lower (depression) threshold, and crossing of the higher (potentiation) threshold, respectively (Fig. 3).

### Further Model Predictions and Future Work

Our simulations predict LTP of CTX to STN synapses after the repetitive delivery of strongly hyperpolarizing inhibitory current pulses to the STN, which should be verified in future experiments (Fig 3g). Under the assumption that similar plasticity mechanisms also act in the GPe, our model further predicts that bursting input to the GPe via the STN, or GPe rebound bursting, may lead to LTP in the GPe. Future experiments in the GPe similar to those by Chu et al. (*28*) and Wang et al. (*53*) could contribute to assessing the validity of this assumption. It has been hypothesized that intra-GPe transmission decorrelates the activity of GPe neurons (*26*), suggesting that LTP in the GPe would counteract abnormal synchrony. Previous studies of a small-scale computational model of the STN-GPe network found that spike-timing-dependent plasticity of GPe to GPe connections alone can lead to the coexistence of synchronized and desynchronized states in this circuit (*64*).

### Modeling Structural Plasticity

In addition to calcium-based synaptic plasticity, we considered calcium-based structural plasticity by implementing a calcium-dependent structural plasticity rule similar to the Butz and Van Ooyen framework (*35*). This structural plasticity mechanism adds or removes synaptic connections based on the difference between the present calcium concentration and a target value, which, in turn, governs firing rates in the stationary state. Detailed knowledge about structural plasticity in the STN-GPe circuit is currently not available; however, widespread synaptic reorganization—in the form of changes in the density of synaptic boutons in the STN-GPe circuit—has been observed in animal models of PD in response to chronic dopamine depletion (*27,29,30,65,66*) or chemogenetic activation of striatal neurons (*29*).

### Cause or Consequence: Synaptic Reorganization in Response to Dopamine Depletion

The observation of synaptic reorganization in the BG in response to chronic dopamine depletion in animal models of PD prompted the discussion of how synaptic reorganization relates to altered neuronal activity in the DD BG (*26*). A critical question in this context is whether synaptic reorganization is the cause, or a consequence, of BG network dysfunction (*26*). To address this question, we reproduced elevated iMSN activity leading to the dominance of the akinetic indirect pathway, a key feature of BG dysfunction according to the classical direct/indirect pathway model (*4–6*). Then, we tested whether this would trigger synaptic reorganization and which plasticity mechanisms may specifically contribute to this synaptic reorganization. In our computational model, the iMSN firing rate increase triggered a series of synaptic changes throughout the entire STN-GPe circuit (Fig. 4). Most of these changes have previously been observed experimentally. Specifically, the loss of CTX to STN connectivity was observed in response to chemogenetically elevated striatal activity (*29*). After chronic dopamine depletion in animal models of PD, experimental studies also found a loss of CTX to STN connectivity (*29, 65, 66*), a gain of GPe to STN synapses (*27*), and a loss of STN to GPe connectivity (*30*).

Our computational model also provides insight into the potential mechanisms underlying synaptic reorganization in the STN-GPe circuit. In our simulations, elevated iMSN activity increased STN firing rates by suppressing inhibitory GPe inputs. Corresponding changes in STN and Gpe firing rates after striatal dopamine depletion have been observed experimentally (*17, 26, 67*). Firing rate changes were followed by an increase in the STN calcium concentrations, because calcium influx through NMDA receptors is related to the product of pre- and post-synaptic firing rates (*68*). Elevations in the STN calcium concentrations were counteracted by structural plasticity; this caused a loss of excitatory CTX to STN connections and a gain of inhibitory GPe to STN connections (Fig. 4g). Despite inhibition by elevated iMSN activity, the GPe calcium concentrations increased due to the elevation in the (pre-synaptic) STN firing rate. The increase was counteracted by structural plasticity by a loss of STN to GPe connections and a gain of GPe to GPe connections.

### Challenging the Existing Neuronal Structural Plasticity Paradigms

The loss of STN to GPe connections observed in this study (Figs. 4i,4i’, and in 6-OHDA mice (*30*)) is difficult to reconcile with the traditional view that structural plasticity aims for homeostasis of neuronal activity, with calcium concentrations acting as a proxy signal for neural firing rates (*35,48,69*). With this traditional view, homeostatic structural plasticity should cause a gain of STN to GPe connections and a loss of GPe to GPe connections to counteract the GPe inhibition by elevated iMSN activity. However, due to our model’s calcium-based structural plasticity mechanism, the mean GPe calcium concentrations increased due to the elevation in pre-synaptic STN activity; this subsequently increased the opening time of post-synaptic NMDA receptors, facilitating calcium influx. This homeostatic mechanism maintains the mean GPe calcium concentrations and results in a loss of STN to GPe connections (as observed experimentally (*30*)) and a gain of GPe to GPe connections, synaptic changes that would be considered anti-homeostatic in the traditional view where structural plasticity is set to balance firing rate deviations from a target value. The gain of GPe to GPe connections has not yet been observed experimentally to date; however, we propose that the increased transmission strength of GPe to GPe connections in 6-OHDA-lesioned rats (*70*) may be mediated by more GPe to GPe synapses and/or a change in the efficacy of existing synapses.

### Other BG Activity Changes in Response to Dopamine Depletion

Besides firing rate modifications, research into animal models of PD revealed a wide range of activity changes in the DD BG. One is pronounced low-frequency bursting of cortical and striatal projection neurons, observed in rodent models of PD (*56, 61*). To test whether synaptic reorganization also occurred in response to cortical and iMSN low-frequency bursting, we modulated cortical and iMSN inputs accordingly. Low-frequency bursting in the cortex and striatum reduced the change in GPe afferent connectivity compared to iMSN firing rate increases alone (compare Fig. 4i and Fig. 4i’). At approximately 25%, the loss of STN to GPe connectivity was closer to the experimentally observed values for GPe-bound connections in 6-OHDA lesioned mice (*30*) than after iMSN firing rate increase. Changes in the number of STN afferents were similar after switching to low-frequency burst input (Fig. 4i’) and increased iMSN firing rate (Fig. 4i). In our computational model, synaptic efficacies were enhanced after switching to low-frequency bursting.

In contrast, they were reduced after switching to increased iMSN firing rates (compare Fig. 4j’ and Fig. 4j). However, the experimental findings regarding changes in synaptic strengths are unclear. Fan et al. suggested that the strengths of GPe to STN synapses may be unchanged following dopamine depletion, based on the observation that the amplitudes of miniature inhibitory postsynaptic currents in the STN were not increased following 6-OHDA lesion (*27*). Chu et al. suggested that the inability of CTX to STN synapses to undergo potentiation following a 6-OHDA lesion in a mouse model may imply that these synapses have been potentiated to their maximum value (*29*).

Our results suggest that synaptic reorganization in the STN-GPe network strongly depends on iMSN and CTX activity characteristics, which vary in different brain states (*17, 19*). In our computational model, structural reorganization was robust concerning input activity changes. In contrast, the dynamics of synaptic efficacies varied based on input activity, likely reflecting input correlations (compare Figs. 4i,j and 4i’,j’). Synaptic efficacy changes can occur on a time scale of minutes (*71*), suggesting that the strengths of individual synapses may depend on the experimental conditions in which recordings are made.

### Exploring the Relationship Between Beta Oscillations and Elevated iMSN Activity

PD is associated with the emergence of exaggerated beta oscillations; however, it is still under debate how such oscillations relate to motor symptoms (*20, 25*). In rat (*17, 19*) and non-human primate models of PD (*18*), a late onset of beta oscillations following chronic dopamine depletion has been observed. This supports the hypothesis that the onset of beta oscillations results from slow adaptation processes triggered by dopamine depletion that increases iMSN activity (*6, 26*). We tested this hypothesis in our computational model by increasing iMSN activity, which results from dopaminergic lesions (*56*). The rapid increase makes it easier to identify the contributions of iMSN activity to firing rates and beta power in the STN and GPe. For example, elevating the iMSN rate leads to an immediate decrease in the GPe firing rate, followed by a further decrease due to structural reorganization (Fig. 4e). The individual contribution of both of these factors would not be as clear if the iMSN firing rate were slowly titrated to its final value.

Even though our model reproduced known synaptic reorganization (Fig. 4), we did not observe a delayed onset of STN and GPe beta oscillations (Fig. 5). This discrepancy might result from slower progression of dopamine depletion within the striatum in experimental studies (*18*), which we did not consider. Furthermore, elevated oscillatory activity was recorded in rats after complete but not after partial 6-OHDA lesion (*72*). The increase in the firing rates of all iMSN in our model might resemble the effects of dopamine depletion of major parts of the striatum or the complete lesion case. It also remains a possibility that upstream changes in synaptic organization, in the motor cortex or striatum, lead to the generation of larger amplitude beta oscillations, which then propagated to the STN and GPe. In general, we find substantial variability in the intensity of beta oscillations depending on cortical and iMSN activity characteristics. This reflects the variety of experimental findings across different animal models and PD patients (*10,20*); in particular, that exaggerated beta oscillations have not been observed in 6-OHDA-lesioned mice (*10*) and that beta oscillations in 6-OHDA rats and MPTP monkeys occur at different frequencies (*20*), where the frequency in rats closer resembles that observed in PD patients. It has been suggested that this variability may result from the involvement of different subcircuits in beta oscillation generation (*25*) and the dependence of abnormal beta oscillations on the brain states (*19*).

### Synaptic and Structural Plasticity Produce Different Changes in Beta Power

While synaptic plasticity changes synaptic efficacies if calcium surpasses specific threshold concentrations, structural plasticity leads to the generation of new and the removal of old synaptic connections in response to deviations from a target concentration. Performing simulations of our computational model without synaptic and/or structural plasticity, we found distinct contributions of either type of plasticity to elevated beta power after input activity changes. Synaptic plasticity alone led to increased beta power minutes after changing input activity (Fig. 5,f), likely caused by the rapid increase of synaptic efficacies during the first minutes of the simulations (Fig. 4h and Fig. 4h’). In contrast, considering only structural plasticity typically reduced beta power, except for STN beta power following the switch to cortical and iMSN low-frequency bursting (Fig. 5c,f). Together, these results demonstrate the essential role of different calcium-based plasticity mechanisms in the generation and/or maintenance of exaggerated beta oscillations in the STN-GPe circuit. They also suggest that synaptic plasticity in response to correlated inputs may further pathological oscillations in the BG and contribute to the brain state dependence of such oscillations, as inputs vary according to the brain state. For instance, in 6-OHDA lesioned rats, beta oscillations were observed in an active state but not during slow-wave sleep states (*19, 67*). Note that a sudden change of iMSN and/or CTX activity led to some degree of elevated beta power, even in the absence of plasticity, indicating that a shift in inputs alone can also lead to elevated beta power in the model.

### Limitations of the Study

Our computational model captures several key aspects of long-term synaptic reorganization within the STN-GPe circuit that have not been previously simulated; however, several limitations exist. Firstly, using single-compartment conductance-based neurons balances biophysical detail and computational efficiency, enabling simulation durations of several days with detailed calcium dynamics (Figs. 4 and 5). Second, our model lacks spatial details of synaptic connectivity and local calcium concentrations through the dendrites, which is suggested to play a vital role in governing heterosynaptic plasticity (*54*). Next, short-term plasticity was not incorporated; however, we acknowledge that this may affect spike patterns such as bursting (*73*). Due to a lack of experimental data, we considered a single-fixed point version of the homeostatic plasticity rule presented by Butz and Van Ooyen (*35, 60*). Even though some experimental evidence supports the existence of a low calcium fixed point for structural plasticity in the rat cerebral cortex (*74*), calcium levels in our simulations never reached such low concentrations.

We also note that the calcium-based plasticity model from Graupner and Brunel approximates the complex secondary messenger signaling system that underlies LTP and LTD (*52*). Our investigation was limited to the STN-GPe circuit; thus, additional network-wide factors were not considered. These include sub-populations of GPe neurons with distinct activity and connectivity patterns (*30, 75*), striato-pallidal loops in which beta activity may originate (*25*), and feedback from the entire cortico-BG circuit. We aim to include this feedback in future versions of the computational model to study its effect on network structure and the onset of exaggerated beta oscillations.

### Conclusions

Our results suggest that homeostatic calcium-based plasticity in the BG may contribute to the observed synaptic reorganization that occurs following chronic DD (*27, 29, 30, 65, 66, 70*). Our computational model generated similar synaptic reorganization in response to elevated iMSN activity, a core tenet of the classical direct/indirect pathway model of BG dysfunction in PD (*4–6*). We also found distinct and partly opposite contributions of synaptic and structural plasticity to the emergence of beta oscillations. Whereas increased synaptic strengths due to synaptic plasticity tended to enhance beta oscillations, structural reorganization in response to elevated iMSN and low-frequency burst input tended to counteract beta oscillations. These results strongly support the hypothesis that synaptic reorganization due to structural plasticity occurs as a corrective mechanism intended to counteract the effects of elevated iMSN input due to striatal dopamine depletion.

## Methods

### Computational Model of Long-Term Plasticity in the STN-GPe Circuit

We developed a computational model of long-term plasticity in the STN-GPe circuit consisting of 250 STN and 750 prototypic GPe neurons simulated using modified conductance-based models (*32, 33*). The STN received 2500 excitatory inputs from the cortex, and the GPe received 7500 inhibitory inputs from the striatum, with each of these inputs simulated using Poisson spike generators. Connections from the cortex to the STN and from STN to GPe are excitatory, and produce AMPA and NMDA post-synaptic currents. Connections from the striatum to the GPe, from GPe to STN, and from GPe to GPe are inhibitory and produce a GABA post-synaptic current (Fig. 1). A calcium-based plasticity rule governs synaptic conductances (*34*). Heterosynaptic plasticity is modeled by specifying that each neuron has three dendritic segments, which possess local calcium concentrations that influence the strengths of all synapses onto that segment (Fig. 6). Each excitatory and inhibitory connection is associated with a randomly chosen segment on the post-synaptic neuron. The number of excitatory and inhibitory connections received by each STN and GPe neuron is controlled by a homeostatic structural plasticity rule that responds to changes in somatic calcium concentrations on a timescale of hours (*35*). Synaptic and structural plasticity of MSN to GPe synapses are not simulated as it has been reported that no change occurs at the synapses between the iMSN and prototypic GPe neurons in 6-OHDA lesioned rodents (*59, 70*).

### Conductance-based neuron models

Single-compartment, point neurons were used to model STN and GPe neurons (*32, 33*). The dynamics of the membrane potential +_8_ of neuron 8 is described by:

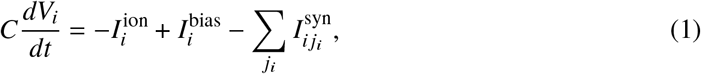

where *C* is the membrane capacitance, 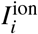 is the sum of all ionic currents, 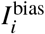 is the applied bias current, and the last term is the sum over all synaptic input currents, where *j*_*i*_ is the index of the *j*_*i*_ th synapse of the postsynaptic neuron *i*. Following, the index *i* refers to quantities associated with the *i*th neuron.

The total ionic current for STN neurons is given by

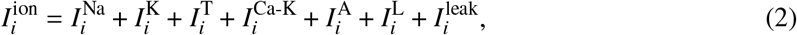

and for GPe neurons it is given by

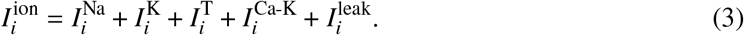

More details on individual ionic currents are detailed in Table 1.

Each ionic current is composed of a conductance *g*^X^, where X specifies the ionic current, a reversal potential *E*^X^, and one or more gating variables G. The conductances are given in Table 2. The reversal potentials are constant for sodium (*E*^Na^ = 40 mV), potassium (*E*^K^ = −90 mV), and leak (*E*^leak^ = −60 mV) currents, whereas the reversal potential for calcium, *E*^Ca^, is derived from the ratio of the external calcium concentration, [Ca^soma^]^out^, to the internal calcium concentration within the soma of neuron *i*, 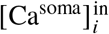, using the Nernst equation:

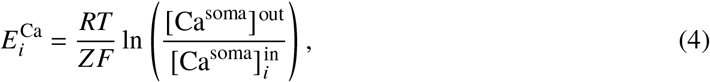

where *R* is the universal gas constant, *T* = 303.15 K is the temperature, *Z* = 2 is the valence of the calcium ions, F is Faraday’s constant, and the external calcium concentration is set to [Ca^soma^]^out^ = 2 ‘M (*32*).

**Table 2:** STN and GPe conductances.

| Conductance | STN (mS/cm<sup>2</sup>) | GPe (mS/cm<sup>2</sup>) |
| --- | --- | --- |
| $g^{\text{Na}}$ | 49 | 40 |
| $g^{\text{K}}$ | 57 | 4.2 |
| $g^{\text{T}}$ | 4.0 | 0.134 |
| $g^{\text{Ca-K}}$ | 1.0 | 0.1 |
| $g^{\text{leak}}$ | 0.35 | 0.04 |
| $g^{\text{A}}$ | 5.0 | |
| $g^{\text{L}}$ | 0.35 | |

The gating kinetics are governed by the equation:

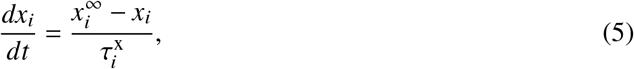

where *x* ∈ {*a, b, c, d*_1_, *d*_2_, *h, m, n, p, q, r*} corresponds to one of the gating variables listed in Supplementary Table 1 and 2.

The steady state (in)activation functions,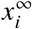, are given by:

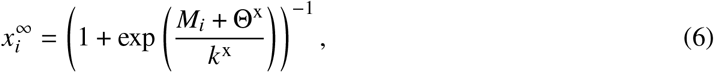

where Θ^X^ are the half (in)activation voltages and *k*^X^ are the half (in)activation slopes. The membrane potential, *M*_*i*_ = *V*_*i*_, governs the response of all gates except *d*_2_ and *r*, which are governed by the internal somatic calcium concentration 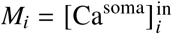.

The fast (in)activation time scales, i.e. *x* ∈ {*a, d*_2_, *m, r*}, are given by:

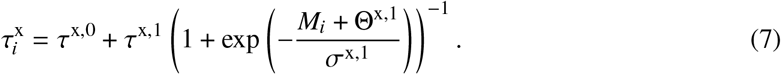

The slow (in)activation time scales, i.e. *x* ∈ {*b, c, d*_1_, *h, n, p, q*}, are given by:

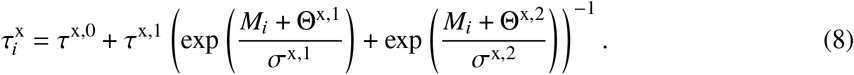

The value for each gating variable may be found in Supplementary Table 1 and 2.

### Somatic and segmental calcium dynamics

The somatic calcium concentration,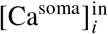, of a given neuron 8 governs the behaviour of calcium-dependent currents via ligand gating, and influences the structural plasticity as described below. The calcium concentration at segment *l*_*i*_ on a neuron *i*,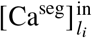, governs the strength of the synapses associated with that segment, via the mechanism described by Graupner and Brunel (*34*). The somatic calcium dynamics are described using the differential equation presented by Otsuka et al. (*32*), which we extended by an additional calcium current arising from the NMDA receptors:

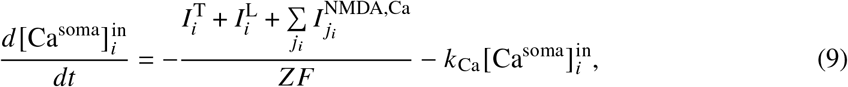

where *k*_Ca_ = 2 ms^-1^ is the calcium removal rate, 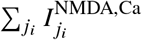 is the calcium influx through all NMDA receptors, where *j*_*i*_ refers to the *j*_*i*_th synapse of neuron *i*. Similarly, the dynamics of the calcium concentrations in the three segments of neuron *i* are described by:

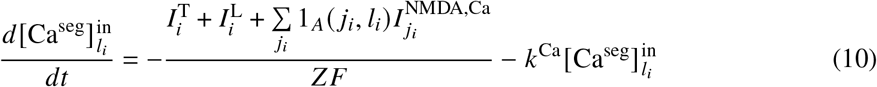

*l*_*i*_ = 0, 1, 2 refers to individual segments of neuron *i* (Fig. 6). The NMDA current will only contribute to the change in calcium concentration if it synapses onto segment *l*_*i*_, as denoted by the indicator function 1_*A*_ (*j*_*i*_, *l*_*i*_) which returns one if synapse *l*_*i*_ connects to segment *l*_*i*_, and zero otherwise.

### Synapses

Synaptic conductances were represented using the model of Rubin et al. (*52*). Ligand gating of currents at the synapse *j* is governed by rise,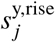, fast, 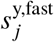, and slow components, 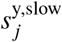:

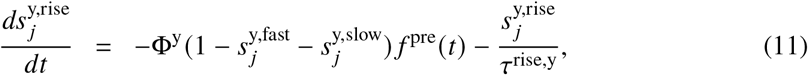

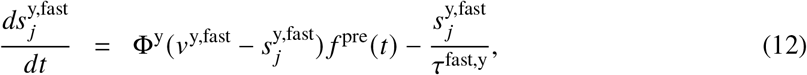

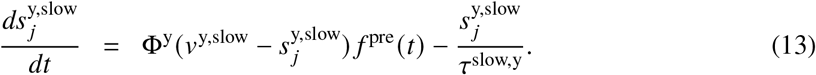

Here, *y* specifies the type of neurotransmitter, i.e., GABA_a_, AMPA, or NMDA. *f* ^pre^ is a step pulse lasting 1 ms, modelling presynaptic transmitter release, which is turned on to activate the postsynaptic synapse, *j*, when a presynaptic spike occurs. *ϕ* is the rate of increase of the gating variables when neurotransmitter is released. *v*^y,slow^ and *v*^y,fast^ are attractor values for the slow and fast gates, and *τ*^rise,y^, *τ*^fast,y^, and *τ*^slow,y^ are the time constants of the rise, fast, and slow components, respectively. The value of each parameter involved in synaptic gating may be found in Table 3.

**Table 3:**
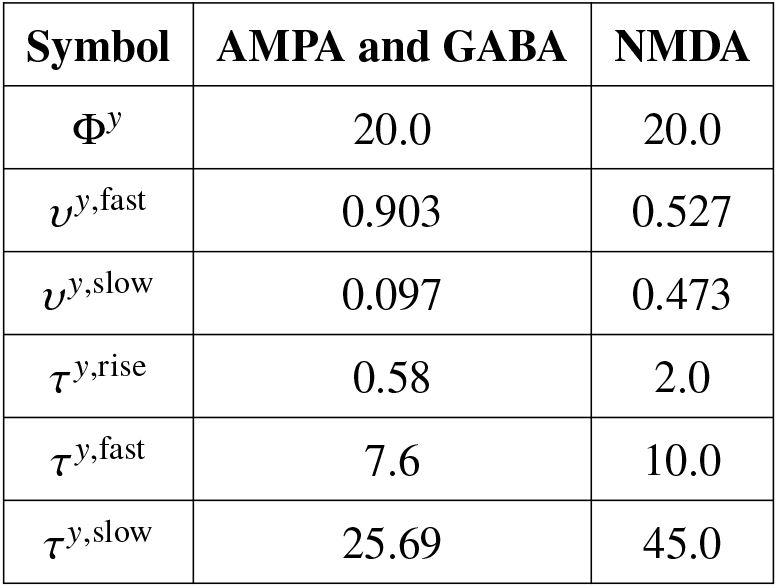
Parameters for AMPA, GABA, and NMDA synapses.

The current from an inhibitory synapse, i.e., *y* = GABA_a_, is given by

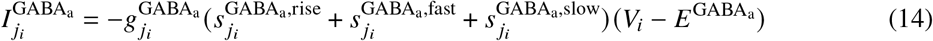

Here, the index *j*_*i*_ refers to the *j*_*i*_ th synapse of the postsynaptic neuron *i*. 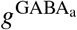 is the maximum conductance,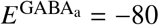 mV the synaptic reversal potential, and *V*_*i*_ the membrane potential of the post-synaptic neuron *i*.

Similarly, the current from the fast AMPA channel of excitatory synapses, with *y* = AMPA, takes the following form:

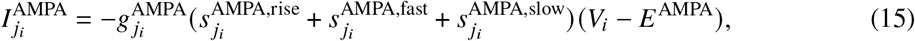

with *g*^AMPA^ being the conductance, and *E*^AMPA^ = 0 mV the reversal potential of the AMPA channels.

In addition, the excitatory synapse model contains NMDA receptors (*y* = NMDA). Current flow through the NMDA receptors is divided into two components, a calcium current, *I*^NMDA,Ca^, and non-specific cationic current, *I*^NMDA,syn^ (*52*):

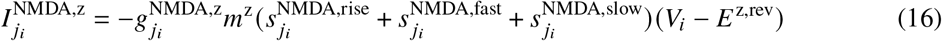

Here, *z* = Ca or *z* = syn. The effect of magnesium block removal on the opening of NMDA receptors is modelled by *m*^z^ with (*52*):

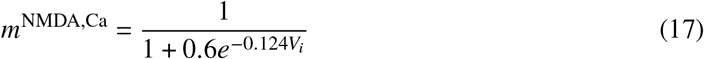

and

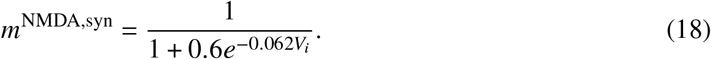

To reproduce the ratios of NMDA to AMPA currents observed in CTX to STN (*28*) and STN to GPe (*30*) synapses, the following relation was assumed *g*^NMDA,syn^ = 1.76^AMPA^. This ensured that the NMDA:AMPA current ratio, recorded under depolarized and hyperpolarized conditions respectively, was 0.25:1.

### Synaptic plasticity

To incorporate synaptic plasticity, we described the individual fast excitatory and inhibitory conductances,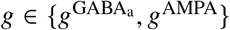, by:

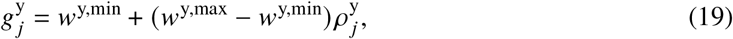

where 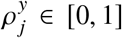 represents the synapse’s efficacy, and *w*^y,min^ and *w*^y,max^ are the minimum and maximum synaptic conductances of synapse type y, respectively. Following, we suppress the index 9 to avoid bulky terms. The dynamics of d^H^ are described using the calcium-based plasticity model presented by Graupner and Brunel (*34*):

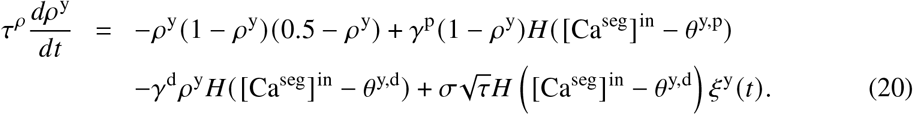

where [Ca^seg^]^in^ is the calcium concentration in the postsynaptic segment. The time scale of calcium-based synaptic plasticity is given by *τ*^*ρ*^ = 30 s, which is in the range suggested by Graupner and Brunel (*34*). The cubic term on the right-hand side of Equation (20), describes the dynamics of 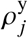 in the absence of calcium fluctuations. In that case, there are two stable fixed points at 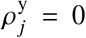 and 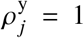 corresponding to down and up states respectively, and an unstable fixed point at 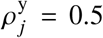. The second term describes synaptic potentiation, scaled by *ρ*^P^. *H*(*x*) is the Heavyside step function that returns a value of one if *x* ≥ 0 and a value of zero otherwise. Thus, the potentiation term contributes to the dynamics of 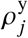 if the segment’s calcium concentration [Ca^seg^]^in^ is greater than the potentiation threshold *θ*^y,p^. The third term describes depotentiation of synapses when the segment’s calcium concentration is above the depression threshold *θ*^y,d^. The final term describes activity-dependent noise. This noise is only present when the calcium concentration is above the depression threshold,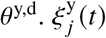 is zero-mean white Gaussian noise, 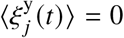 and 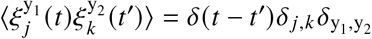. Here, *δ*(*x*) is the Dirac delta distribution and *δ*_*a,b*_ = 1 if *a* = *b* and zero otherwise.

### Structural plasticity

We incorporated a model of homeostatic, calcium-based, activity-dependent structural plasticity based on the linear Butz and Van Ooyen algorithm (*35,60*) into the neurons of the STN and GPe. The original algorithm governed synaptic connections between excitatory neurons, and was modified here to incorporate inhibitory synapses. Each neuron contains a number of axonal and dendritic elements, which can merge with their opposing element on another neuron to form synapses. Excitatory axonal elements may merge with available excitatory dendritic elements to form new excitatory synapses, and likewise available inhibitory axonal and dendritic elements may merge to form inhibitory synapses. The number of excitatory dendritic elements, 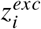, and inhibitory dendritic elements,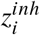, available within a given neuron evolves based on the dynamics of the long-term average calcium concentration 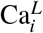. The dynamics of 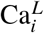 are described as a simple low-pass filter of the somatic calcium concentration:

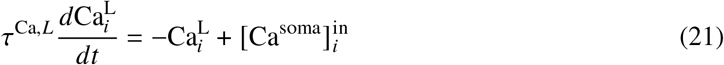

Here, *τ*^Ca,L^ = 10 s is the slow time scale at which the calcium concentration is averaged. Unlike in the Butz and van Ooyen model, the numbers of excitatory,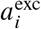, and inhibitory axonal elements,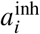, remain static at a large value 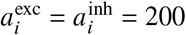. Thus, the formation of synapses depends only on the somatic calcium concentration in the post-synaptic neuron. The change in the number of available excitatory or inhibitory dendritic elements is a linear function of 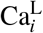, given by:

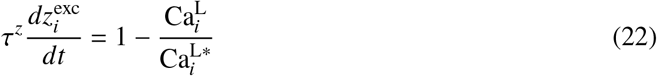

and

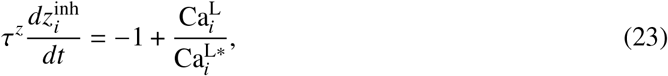

respectively.

The time constant for element growth is set to *τ*^z^ = 3600 s, and 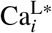 is the target value for 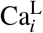. Homeostasis of 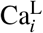 occurs as a result of the relationships established in the above equations, such that when 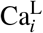 falls below 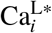, excitatory connections are added and inhibitory connections are lost, and vice versa when 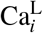 is greater than 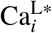. The number of vacant excitatory 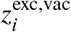 or inhibitory elements 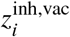 is the floor of the difference between the number of available and the number of bound elements 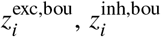, given by: 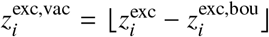, or 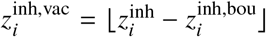. Synapses may form when vacant axonal and dendritic elements are available. While the variables evolved continuously at each timestep, the updates in connectivity were simulated to occur every 200 ms. At each connectivity update, a connection could be formed between a vacant dendritic element and a randomly chosen axonal element, with probability 0.1.

### Cortical and iMSN inputs

The activity of individual cortical and iMSN afferents projecting to the STN and GPe, respectively, were modelled by Poisson spike generators. We considered two types of spike generators: spiking and pause-bursting ones. Spiking afferent inputs were modelled as Poisson spike trains, with constant rate parameter 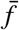. Pause-bursting activity was realized by switching between a high rate (bursting), a low rate (spiking) and no activity (pause).

To model different input activity patterns, we varied the rate parameter for spiking afferents,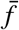, as well as the portion of bursting afferents, *P*^b^, and the parameters characterizing their activity pattern.

Bursting occurred at an average frequency of *F*^bo^ = 1/*d*^bo^, where the time between subsequent burst onsets, *t*^b^, was drawn from a normal distribution with mean *d*^bo^ and a standard deviation of 0.1 s. During bursts, *n*^b^ spikes with exponentially distributed interspike intervals were generated with mean interval 1/ *f* ^b^ and Poisson-distributed *n*^b^ with *λ*= *f* ^b^*d*^b^, where *d*^b^ measures the burst duration. After the last spike, the afferent switched to spiking mode with rate parameter *f* ^s^ until *t*^b^ - *d*^p^ with 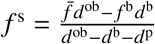, ensuring an overall mean firing rate of 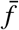. Parameters for the different input activity patterns are given in Table 4.

**Table 4:**
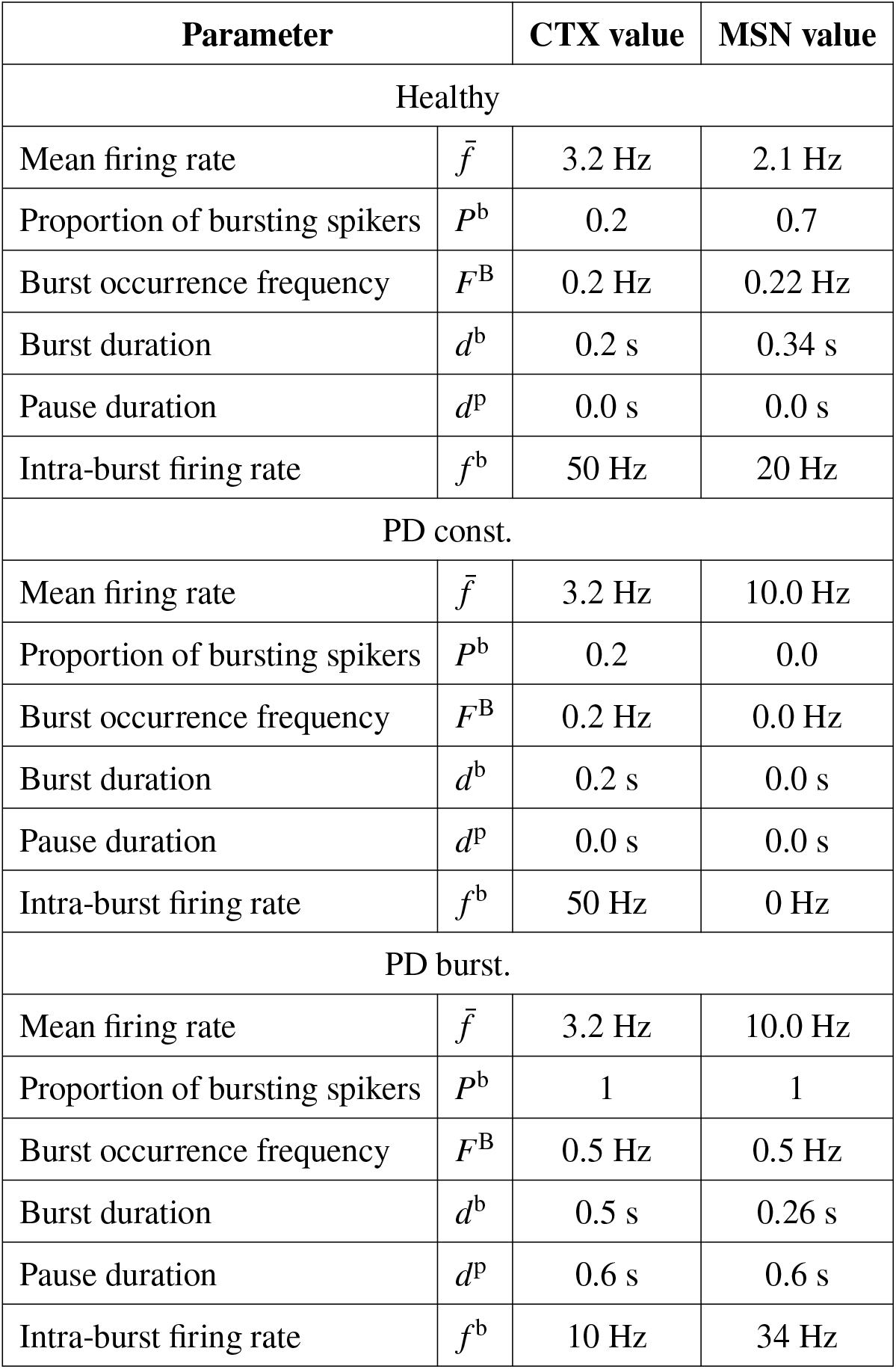
Activity parameters for spike generators under different conditions. Values were taken where appropriate from the available literature (*56, 61*). Here, “PD const.” and “PD burst.” refer to PD states with fixed iMSN and cortical firing rates and time-dependent iMSN and cortical firing rates mimicking low-frequency bursting, respectively (see also Fig. 2).

| Parameter |  | CTX value | MSN value |
| --- | --- | --- | --- |
| Healthy |  |  |  |
| Mean firing rate | $\bar{f}$ | 3.2 Hz | 2.1 Hz |
| Proportion of bursting spikers | $P^b$ | 0.2 | 0.7 |
| Burst occurrence frequency | $F^B$ | 0.2 Hz | 0.22 Hz |
| Burst duration | $d^b$ | 0.2 s | 0.34 s |
| Pause duration | $d^p$ | 0.0 s | 0.0 s |
| Intra-burst firing rate | $f^b$ | 50 Hz | 20 Hz |
| PD const. |  |  |  |
| Mean firing rate | $\bar{f}$ | 3.2 Hz | 10.0 Hz |
| Proportion of bursting spikers | $P^b$ | 0.2 | 0.0 |
| Burst occurrence frequency | $F^B$ | 0.2 Hz | 0.0 Hz |
| Burst duration | $d^b$ | 0.2 s | 0.0 s |
| Pause duration | $d^p$ | 0.0 s | 0.0 s |
| Intra-burst firing rate | $f^b$ | 50 Hz | 0 Hz |
| PD burst. |  |  |  |
| Mean firing rate | $\bar{f}$ | 3.2 Hz | 10.0 Hz |
| Proportion of bursting spikers | $P^b$ | 1 | 1 |
| Burst occurrence frequency | $F^B$ | 0.5 Hz | 0.5 Hz |
| Burst duration | $d^b$ | 0.5 s | 0.26 s |
| Pause duration | $d^p$ | 0.6 s | 0.6 s |
| Intra-burst firing rate | $f^b$ | 10 Hz | 34 Hz |

### Model fitting procedure

The model, in its default state, was tuned to reflect the dopamine-intact STN-GPe circuit of the rat, for which the neural firing patterns and rates are well-described in the literature (*55, 56*). The synaptic weights giving rise to this behaviour are detailed in Table 5.

**Table 5:**
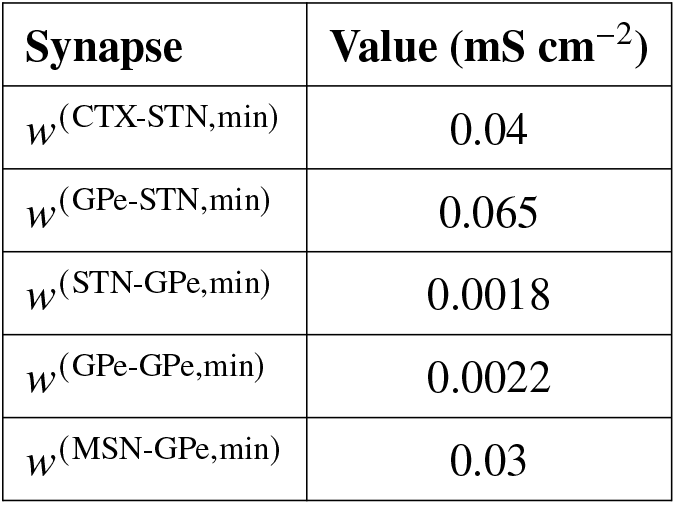
Synaptic conductances and their minimal values.

| Synapse | Value (mS cm<sup>-2</sup>) |
| --- | --- |
| $w^{(\text{CTX-STN},\text{min})}$ | 0.04 |
| $w^{(\text{GPe-STN},\text{min})}$ | 0.065 |
| $w^{(\text{STN-GPe},\text{min})}$ | 0.0018 |
| $w^{(\text{GPe-GPe},\text{min})}$ | 0.0022 |
| $w^{(\text{MSN-GPe},\text{min})}$ | 0.03 |

### Synaptic plasticity

The Graupner and Brunel (2012) plasticity model (*34*) has four free parameters per synapse *γ*^SYN,p^, *γ*^SYN,d^, *θ*^SYN,p^, and *θ*^SYN,d^, and our computational model contains four distinct types of plastic synapses SYN ∈ {CS, GS, SG, GG}. It has been observed experimentally that plasticity at CTX to STN and GPe to STN synapses is heterosynaptic: activity at CTX to STN synapses can influence the strength of GPe to STN synapses (*28*). For simplicity, we assumed that the same mechanism is present between STN to GPe and GPe to GPe synapses. Although no experimental evidence is available to prove or disprove this, heterosynaptic plasticity is observed in many neuron types (*54*). To model this heterosynaptic effect, the potentiation, *θ*^SYN,p^, and depression thresholds, *θ*^SYN,d^ were set equal for excitatory and inhibitory synapses on each neuron. This results in a single potentiation threshold for the STN *θ*^STN,p^ = *θ*^CS,p^ = *θ*^GS,p^, and for the GPe *θ*^GPe,p^ = *θ*^SG,p^ = *θ*^GG,p^, and a single depression threshold for the STN *θ*^STN,d^ = *θ*^CS,d^ = *θ*^GS,d^ and for the GPe *θ*^GPe,d^ = *θ*^SG,d^ = *θ*^GG,d^. An additional constraint necessary to reproduce plasticity behaviour observed in the STN is that potentiation occurs at a higher calcium level compared to depression (*53*), so *θ*^STN,p^ > *θ*^S STN,d^, and similarly it was assumed that *θ*^GPe,p^ > *θ*^GPe,d^. For simplicity, the same potentiation and depression rates were used for all synapses, *γ*^p^ = *γ*^CS,p^ = *γ*^GS,p^ = *γ*^SG,p^ = *γ*^GG,p^ and *γ*^d^ = *γ*^CS,d^ = *γ*^GS,d^ = *γ*^SG,d^ = *γ*^GG,d^. Potentiation and depression thresholds were determined through an iterative trial and error process to balance long-term stability and responsiveness to excitatory inputs. Plasticity rules based on correlations between pre- and post-synaptic activity are inherently unstable as they induce a positive feedback loop whereby correlated activity at a synapse strengthens that synapse, increasing the likelihood that future activity will be correlated and leading to a state where all synaptic efficacies are at their maximum value (*76*). In the Graupner and Brunel model, the trade-off between sensitivity to correlations and stability of *ρ*^SYN^ can be controlled by adjusting *θ*^SYN,p^, *θ*^SYN,d^, and *γ*^p^. The depression rate can be used with the plasticity time constant to control the rate at which plastic changes occur. If the plasticity is too sensitive, then all weights will move towards their attractor at *ρ*^SYN^ = 1. Conversely, if the plasticity is too stable, then it will not respond to any increases in activity or correlation among presynaptic neurons. As observed in Chu et al. (2015) (*28*), a bursting protocol consisting of 300 ms of 50 Hz firing repeated at Hz, among all presynaptic cortical afferents, should potentiate CTX to STN and GPe to STN synapses to double their initial conductance. No similar protocols were found in the literature to quantify STN to GPe or GPe to GPe efficacy, so it was assumed that this protocol would result in potentiation of GPe afferents. Future experiments applying such protocols to the GPe could yield information on whether this assumption is valid. It was desired that the mean value of *ρ*^SYN^ for each synapse should be close to 0.33, to reproduce the effects observed in experiments for the STN where conductance of afferent cortical and pallidal synapses could be reduced by 50%, which can occur when *ρ*^SYN^ = 0, or increased by 100%, which can occur when *ρ*^*SYN*^ = 1 (*28, 53*). Initial values for potentiation and depression thresholds were chosen based on the distribution of calcium concentrations within each segment; these values were then adjusted, along with the potentiation and depression rates, to achieve stable and responsive plasticity. The set of parameters defining the plasticity mechanism may be found in Table 6.

**Table 6:** Synaptic and structural plasticity parameters.

| Symbol | Value |
| --- | --- |
| $\theta^{(\text{STN},p)}$ | 17.8 nM |
| $\theta^{(\text{STN},d)}$ | 12.0 nM |
| $\theta^{(\text{GPe},p)}$ | 0.29 nM |
| $\theta^{(\text{GPe},d)}$ | 0.2 nM |
| $\gamma^p$ | 200 s<sup>-1</sup> |
| $\gamma^d$ | 20 s<sup>-1</sup> |
| $\text{Ca}^{(\text{STN},L^*)}$ | 3.86 nM |
| $\text{Ca}^{(\text{GPe},L^*)}$ | 0.058 nM |
| $\tau^{(\text{STN},z)}$ | 300 s |
| $\tau^{(\text{GPe},z)}$ | 300 s |

### Structural Plasticity

The linear Butz and Van Ooyen model has two parameters governing the structural plasticity in each neuron, the target calcium concentrations Ca^STN,L*^ and Ca^GPe,L*^, and the time constants for dendritic element growth *τ*^STN,z^ and *τ*^GPe,z^. The time constants were set equal *τ*^STN,z^ = *τ*^GPe,z^ = 1800 s, and the target calcium concentrations were set to the steady state value determined in simulations with structural plasticity disabled. These values are reported in Table 6.

### Simulation details

The model was implemented using C++ and the GPU parallelization library CUDA. A CUDA kernel was written to solve the system of equations governing the conductance-based neurons and the synapse dynamics. This allowed the system to be solved in parallel, thus significantly decreasing the computational time required for large, long-duration simulations. The conductance-based models are solved using the “Strang splitting” method, which has been demonstrated to preserve the correct limit cycle behavior in Hodgkin-Huxley type neurons while allowing the use of larger timestep (*77*), and the synaptic equations were solved using the forward Euler method. To reduce the computational overhead associated with sending data to and from the GPU, a large number of iterations were solved continuously before control was returned to the CPU. Typically, 6400 iterations were used, corresponding to 200 ms of simulated time with the default timestep of 0.03125 ms. CPU-bound code was responsible for initializing the simulation, generating stochastic spikes of cortical and striatal afferents, performing the updates to network connectivity that arise from the structural plasticity algorithm, and saving results. Custom scripts were written in Python to manage the model’s initialization and running, as well as perform data processing. Simulations typically required 3.5 seconds of real time for one second of simulated time using an Nvidia 3060ti GPU (Nvidia Corporation, Santa Clara, CA) and an Intel i7-9700k CPU (Intel Corporation, Santa Clara, CA). Simulations were run on the UCD Sonic high-performance computing cluster, and Stanford University’s Sherlock computing cluster. Further simulations were run on a local machine with an NVIDIA-sponsored Titan V GPU and Intel Core i9-10980XE CPU with 3.00 GHz.

Networks were initialized as follows: The average in-degree (number of incoming connections per post-synaptic neuron) of each synapse type was initialized to 10. Pre- and postsynaptic neurons were paired randomly according to a uniform distribution until this average in-degree was met. Membrane potentials were initialized between −80 mV and 50 mV, and gating variables for neurons and synapses were initialized between zero and one using a uniform random distribution. To generate stationary networks, we ran simulations for 13 h of simulated time, starting with the default cortical and iMSN activity described in Table 4 (Healthy). Corresponding data for the last hour of these simulations are shown as pre-input change conditions in Figs. 4, and 5. Then, instantaneous changes of cortical and iMSN input were performed, and simulations were continued for the simulated times shown in Figs. 4, and 5.

## Data evaluation

### Beta Power Calculation

The instantaneous population firing rates were estimated by binning spike times into 1 ms intervals and averaging the number of spikes across the population at each interval. The power spectrum of the mean population spike train was then calculated using Welch’s method, with a window size of 1024 samples, and 50% window overlap for spike train samples lasting 60 seconds. The integral beta power was estimated by integrating the power spectral density between 13 and 30 Hz.

### Statistics

We ran simulations for ten random realizations of initial conditions for voltage, gating, and synaptic variables, as well as synaptic connectivity, noise, and cortical and iMSN (inhomogeneous) Poisson inputs, for each condition. Figs. 4 and 5 show the averages of these ten simulations (thick-colored curves) and the results of individual simulations (thin-colored curves).

### Research Standards

The parameters of our computational model were tuned to reflect the firing rates and neural activity observed in the dopamine-intact STN-GPe circuit of the rat (*55, 56, 78–80*). Synaptic and structural plasticity mechanisms were selected and parameterized in order to reproduce long term dynamics that have been observed in animal models of PD (*27, 28, 30, 53, 65*). The “Model Fitting Procedure” section of the Methods describes the process in more detail.

## Author contributions

Conceptualization: C.M., J.A.K., M.L., P.A.T.; Data curation: C.M., J.A.K.; Formal analysis: C.M., J.A.K.; Investigation: C.M., J.A.K., M.L., P.A.T.; Software: C.M.; Methodology: C.M., J.A.K., M.L., P.A.T.; Validation: C.M., J.A.K., M.L., P.A.T.; Visualization: C.M., J.A.K.; Writing – original draft: C.M., J.A.K.; Writing – review & editing: C.M., J.A.K., M.L., P.A.T.; Funding acquisition: C.M., M.L., P.A.T.; Project administration: M.L., P.A.T.; Resources: M.L., P.A.T.; Supervision: J.A.K., M.L., P.A.T.

## Data Availability Statement

The code that was used to generate the figures in this article will be made available on publication (url is added on publication). Further questions can be directed to the Corresponding Author.

