## Supplementary Materials - Tables 1-2 for "Calcium-Based Synaptic and Structural Plasticity Link Pathological Activity to Synaptic Reorganization in Parkinson’s Disease"

Tables S1, S2

**Table S1.**
STN gating parameters

| Parameter | Values | Parameter | Values | Parameter | Values |
|---|---|---|---|---|---|
| $\boldsymbol{C_m}$ | 1 μF/cm$^2$ | $\theta_r$ | 0.17e-3 mM | $\tau_r$ | 2 ms |
| $\boldsymbol{L}$ | 60 μm | $k_a$ | -14.7 mV | $\theta_a^\tau$ | -40 mV |
| $\boldsymbol{Diam}$ | 60 μm | $k_b$ | 7.5 mV | $\theta_b^{\tau 1}, \theta_b^{\tau 2}$ | -60, -40 mV |
| $\boldsymbol{g_l}$ | 0.35 mS/cm$^2$ | $k_c$ | -5 mV | $\theta_c^{\tau 1}, \theta_c^{\tau 2}$ | -27, 50 mV |
| $\boldsymbol{g_{Na}}$ | 49 mS/cm$^2$ | $k_{d_1}$ | 7.5 mV | $\theta_{d_1}^{\tau 1}, \theta_{d_1}^{\tau 2}$ | -40, -20 mV |
| $\boldsymbol{g_K}$ | 57 mS/cm$^2$ | $k_{d_2}$ | 0.02 μM | $\theta_h^{\tau 1}, \theta_h^{\tau 2}$ | -50, -50 mV |
| $\boldsymbol{g_A}$ | 5 mS/cm$^2$ | $k_h$ | 6.4 mV | $\theta_m^\tau$ | -53 mV |
| $\boldsymbol{g_L}$ | 15 mS/cm$^2$ | $k_m$ | -8 mV | $\theta_n^{\tau 1}, \theta_n^{\tau 2}$ | -40, -40 mV |
| $\boldsymbol{g_T}$ | 5 mS/cm$^2$ | $k_n$ | -14 mV | $\theta_p^{\tau 1}, \theta_p^{\tau 2}$ | -27, -102 mV |
| $\boldsymbol{g_{Ca-K}}$ | 1 mS/cm$^2$ | $k_p$ | -6.7 mV | $\theta_q^{\tau 1}, \theta_q^{\tau 2}$ | -50, -50 mV |
| $\boldsymbol{E_l}$ | -60 mV | $k_q$ | 5.8 mV | $\sigma_a$ | -0.5 mV |
| $\boldsymbol{\theta_a}$ | -45 mV | $k_r$ | -0.08 μM | $\sigma_b^1, \sigma_b^2$ | -30, 10 mV |
| $\boldsymbol{\theta_b}$ | -90 mV | $\tau_a^0, \tau_a^1$ | 1, 1 ms | $\sigma_c^1, \sigma_c^2$ | -20, 15 mV |
| $\boldsymbol{\theta_c}$ | -30.6 mV | $\tau_b^0, \tau_b^1$ | 0, 200 ms | $\sigma_{d_1}^1, \sigma_{d_1}^2$ | -15, 20 mV |
| $\boldsymbol{\theta_{d_1}}$ | -60 mV | $\tau_c^0, \tau_c^1$ | 45, 10 ms | $\sigma_h^1, \sigma_h^2$ | -15, 16 mV |
| $\boldsymbol{\theta_{d_2}}$ | 0.1 μM | $\tau_{d_1}^0, \tau_{d_1}^1$ | 400, 500 ms | $\sigma_m$ | -0.7 mV |
| $\boldsymbol{\theta_h}$ | -45.5 mV | $\tau_h^0, \tau_h^1$ | 0, 24.5 ms | $\sigma_n^1, \sigma_n^2$ | -40, 50 mV |
| $\boldsymbol{\theta_m}$ | -40 mV | $\tau_m^0, \tau_m^1$ | 0.2, 3 ms | $\sigma_p^1, \sigma_p^2$ | -10, 15 mV |
| $\boldsymbol{\theta_n}$ | -41 mV | $\tau_n^0, \tau_n^1$ | 0, 11 ms | $\sigma_q^1, \sigma_q^2$ | -15, 16 mV |
| $\boldsymbol{\theta_p}$ | -56 mV | $\tau_p^0, \tau_p^1$ | 5, 0.33 ms | $I_{bias}$ | -1.0 nA |
| $\boldsymbol{\theta_q}$ | -85 mV | $\tau_q^0, \tau_q^1$ | 0, 400 ms | | |

**Table S2.**
GPe gating parameters

| Parameter | Values | Parameter | Values | Parameter | Values |
|---|---|---|---|---|---|
| $\boldsymbol{C_m}$ | 1 µF/cm$^2$ | $\theta_r$ | 0.17e-3 mM | $\tau_r$ | 2 ms |
| $\boldsymbol{L}$ | 60 µm | $k_a$ | -14.7 mV | $\theta_a^\tau$ | -40 mV |
| $\boldsymbol{Diam}$ | 60 µm | $k_b$ | 7.5 mV | $\theta_b^{\tau 1}, \theta_b^{\tau 2}$ | -60, -40 mV |
| $\boldsymbol{g_l}$ | 0.35 mS/cm$^2$ | $k_c$ | -5 mV | $\theta_c^{\tau 1}, \theta_c^{\tau 2}$ | -27, 50 mV |
| $\boldsymbol{g_{Na}}$ | 49 mS/cm$^2$ | $k_{d_1}$ | 7.5 mV | $\theta_{d_1}^{\tau 1}, \theta_{d_1}^{\tau 2}$ | -40, -20 mV |
| $\boldsymbol{g_K}$ | 57 mS/cm$^2$ | $k_{d_2}$ | 0.02 µM | $\theta_h^{\tau 1}, \theta_h^{\tau 2}$ | -50, -50 mV |
| $\boldsymbol{g_A}$ | 5 mS/cm$^2$ | $k_h$ | 6.4 mV | $\theta_m^\tau$ | -53 mV |
| $\boldsymbol{g_L}$ | 15 mS/cm$^2$ | $k_m$ | -8 mV | $\theta_n^{\tau 1}, \theta_n^{\tau 2}$ | -40, -40 mV |
| $\boldsymbol{g_T}$ | 5 mS/cm$^2$ | $k_n$ | -14 mV | $\theta_p^{\tau 1}, \theta_p^{\tau 2}$ | -27, -102 mV |
| $\boldsymbol{g_{Ca-K}}$ | 1 mS/cm$^2$ | $k_p$ | -6.7 mV | $\theta_q^{\tau 1}, \theta_q^{\tau 2}$ | -50, -50 mV |
| $\boldsymbol{E_l}$ | -60 mV | $k_q$ | 5.8 mV | $\sigma_a$ | -0.5 mV |
| $\boldsymbol{\theta_a}$ | -45 mV | $k_r$ | -0.08 µM | $\sigma_b^1, \sigma_b^2$ | -30, 10 mV |
| $\boldsymbol{\theta_b}$ | -90 mV | $\tau_a^0, \tau_a^1$ | 1, 1 ms | $\sigma_c^1, \sigma_c^2$ | -20, 15 mV |
| $\boldsymbol{\theta_c}$ | -30.6 mV | $\tau_b^0, \tau_b^1$ | 0, 200 ms | $\sigma_{d_1}^1, \sigma_{d_1}^2$ | -15, 20 mV |
| $\boldsymbol{\theta_{d_1}}$ | -60 mV | $\tau_c^0, \tau_c^1$ | 45, 10 ms | $\sigma_h^1, \sigma_h^2$ | -15, 16 mV |
| $\boldsymbol{\theta_{d_2}}$ | 0.1 µM | $\tau_{d_1}^0, \tau_{d_1}^1$ | 400, 500 ms | $\sigma_m$ | -0.7 mV |
| $\boldsymbol{\theta_h}$ | -45.5 mV | $\tau_h^0, \tau_h^1$ | 0, 24.5 ms | $\sigma_n^1, \sigma_n^2$ | -40, 50 mV |
| $\boldsymbol{\theta_m}$ | -40 mV | $\tau_m^0, \tau_m^1$ | 0.2, 3 ms | $\sigma_p^1, \sigma_p^2$ | -10, 15 mV |
| $\boldsymbol{\theta_n}$ | -41 mV | $\tau_n^0, \tau_n^1$ | 0, 11 ms | $\sigma_q^1, \sigma_q^2$ | -15, 16 mV |
| $\boldsymbol{\theta_p}$ | -56 mV | $\tau_p^0, \tau_p^1$ | 5, 0.33 ms | $I_{bias}$ | -1.0 nA |
| $\boldsymbol{\theta_q}$ | -85 mV | $\tau_q^0, \tau_q^1$ | 0, 400 ms | | |
